# Buckling instability underlies vertebral segmentation during axolotl tail regeneration

**DOI:** 10.1101/2024.01.31.577464

**Authors:** Wouter Masselink, Tobias Gerber, Vijayishwer Singh Jamwal, Francisco Falcon, Tom Deshayes, Riley Grindle, Ryan P. Seaman, Marco Röcklinger, Sofia-Christina Papadopoulos, Gisela Deneke, Adrija Adhikary, Orestis G. Andriotis, Marko Pende, Yuka Taniguchi-Sugiura, Tzi-Yang Lin, Thomas Kurth, Jingkui Wang, Detlev Arendt, Ji-Feng Fei, Barbara Treutlein, Fred W. Kolling, Philipp J. Thurner, Joel H. Graber, Edouard Hannezo, Elly M. Tanaka, Prayag Murawala

## Abstract

Primary body-axis development is a highly conserved process that proceeds through somitogenesis and subsequent subdivision into dermatome, myotome, and sclerotome. Defects in somitic-clock genes such as *Hes7* lead to vertebral-segmentation defects in mice and fish. Here we show that in the axolotl, although *Hes7* is necessary for proper embryonic vertebral segmentation, it is— surprisingly—dispensable during tail regeneration. We investigated the mechanism of vertebral segmentation during regeneration which initially occurs through extension of a cartilage rod ventral to the spinal cord. We find that the regenerating cartilage rod undergoes a periodic wrinkling that provides a template for vertebral segmentation. Via direct mechanical measurements and biophysical perturbations, we show that a model of compression-induced buckling instability can predict vertebral segmentation. The cartilage rod and other somitic derivatives (muscle, cartilage, tendon, fibroblasts) arise from tendon-like, *Lfng^+^* multi-potent mesenchymal progenitors, which display a gene regulatory state distinct from somitic progenitors. In summary, we uncover a mechanism of vertebral segmentation during axolotl tail regeneration that is distinct from the somite-based developmental mechanism.

## Main text

Regeneration of the axolotl tail, which is composed of a spinal cord surrounded by segmented vertebrae and muscle, is a rare example of primary body-axis regeneration among vertebrates. While the cellular and molecular basis of spinal-cord regeneration in axolotl has been characterized^1,2^, very little is known about how somitic derivatives such as vertebrae are regenerated in axolotl. In vertebrate animals, embryonic development and regeneration have similar outcomes, but their starting points differ, and the question of when the pathways underlying regeneration converge with those underlying development has fascinated generations of researchers^3^. Beyond the differences in starting points, patterning during regeneration of post-embryonic structures occurs at much larger spatial scales compared to embryonic patterning, which suggests that different mechanisms could be at play. During vertebrate embryonic development, patterning of the primary body axis is dependent on somitogenesis. Somites are epithelial balls of cells on either side of the notochord and are specified through the clock-and- wavefront model^4,5^. Somites contain multipotent progenitors that respond to cues from the notochord and then lay out the architecture of the tail; these progenitors are the source of vertebrae and myomeres in addition to other connective-tissue cell types^6,7^. In the axolotl, regeneration takes place in the absence of the notochord and somites; instead, blastema cells condense into a cartilage rod, after which vertebrae emerge^8^. In this study we asked: what is the cell source and mechanism of vertebral patterning during regeneration?

## Hallmarks of somitogenesis are absent during tail regeneration

We investigated two hallmarks of vertebral segmentation during embryogenesis and regeneration, i.e., the formation of epithelial somites, and their subsequent resegmentation. During embryogenesis in axolotls and other vertebrates, the tail contains histologically distinct somites that give rise to vertebrae, muscles, and other types of connective-tissue cells. We found that during regeneration of the axolotl tail, the tail blastema does not contain histologically distinct somites, in contrast to embryogenesis (Fig. 1a, b, Extended Data Fig. 1a). Instead, the earliest visibly segmented structures during regeneration are myomeres which appear around 14 days post amputation (dpa) (Fig.1b, Extended Data Fig. 1a). Nevertheless, despite the absence of somites, both the number of myomeres and vertebrae are correctly re-established in the axolotl tail (Fig. 1c-e) (uninjured: 20.10 +/- 1.45 vs regenerated 19.20 +/- 2.10) ^9^. Axolotl embryos, similar to many other vertebrates, undergo resegmentation, where rostro-caudal specification of somites into *Uncx^+^*and *Tbx18^+^* domains results in a half-segment off-set between myomeres and vertebrae (Fig. 1f-h)^10–12^. Consistent with our observation of a histological absence of somites during regeneration (Fig. 1b, Extended Data Fig. 1a), we found that somite-dependent resegmentation is also absent during regeneration (Fig. 1i-l).

**Figure 1:**
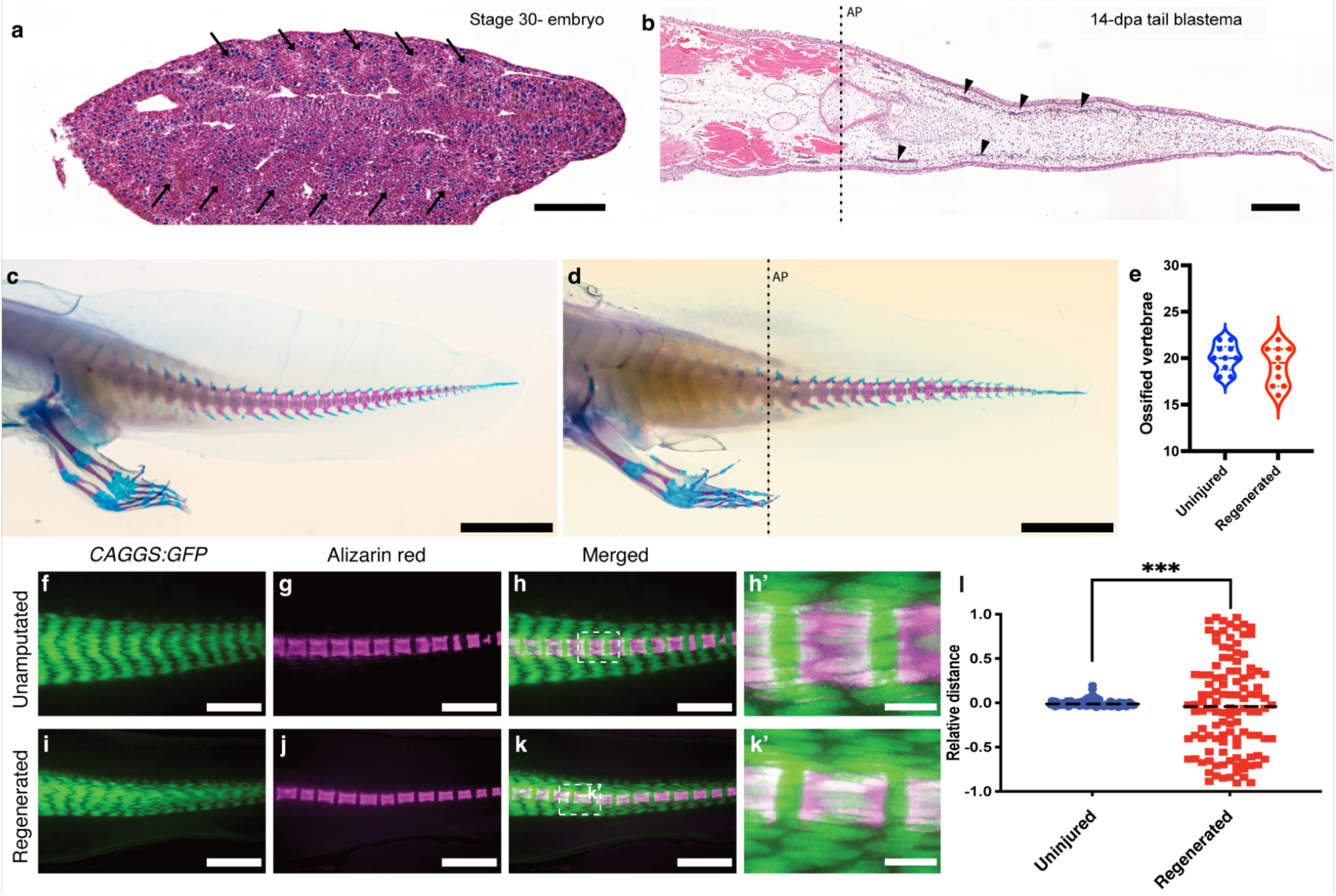
Characterization of axolotl tail regeneration. **a-b,** Coronal section of a H&E-stained stage-30 axolotl embryo (a) and a 14-dpa tail blastema (b). Arrows indicate epithelialized somites (a), and arrowheads indicate segmented myomeres (b). **c-d,** Vertebral segmentation of the uninjured tail (c) is re-established in stage-matched 5-month regenerated tail (d), dashed line indicates amputation plane. **e,** Quantification of ossified vertebrae numbers posterior to the cloaca in uninjured (c) and stage-matched 5-month regenerated (d) animals (n= 10). **f-k**, Stereoscopic images of a *CAGGS:GFP* transgenic axolotl labeled with Alizarin red S in uninjured (f-h) and regenerated (i-k) tail. **l,** Quantification of muscle vertebrae off-set in the tail of h and k (n=112) (off-set is calculated based on the shortest distance from the middle of a vertebra to the nearest myomere boundary. Statistical analysis e: MZ-test. ns: not significant. l: F-test. ns: not significant, ** p< 0.01, *** p< 0.001. Dashed lines indicate amputation plane (ap). Scale bars: a: 200 μm, b: 500 μm, c-d: 1 cm. f-k: 1000 μm. h’, k’: 200 μm.

To understand the processes underlying re-establishment of periodic segments during axolotl tail regeneration, we performed Xenium spatial transcriptomics (Fig. 2a) on stage-28 embryos and on tail blastemas at 3, 7, 10, and 14 dpa (Fig. 2b-e, Extended Data Fig. 1b-i, 2). We selected 100 target genes to visualize not only cell and lineage identity but also pathways known to participate in tissue patterning and signaling, including distal HOX genes, and the FGF, Notch, retinoic acid (RA), Wnt, BMP, and SHH signaling pathways (Extended Data Table 1). Using Xenium clustering analysis, we identified eight cell types in embryo and eight in tail blastema (Fig. 2c, e, Extended Data Fig. 2a, h, o).We found that both embryos and blastemas express distal HOX genes, *Fgfr1*, *Wnt5a,* and the Wnt/Β-cat downstream target *Aldh1a2*, as well as the cell identity markers *Scx, Lfng,* and *Meox1*(Extended Data Fig. 1b-i, 2c-e, j-l, q-s). While *Hes7*, *Tbx6*, and *Mesp2* expression was present in the embryonic tail bud, it was absent in the tail blastema (Fig. 2f, g, Extended Data Fig. 2f, m, t).

**Figure 2:**
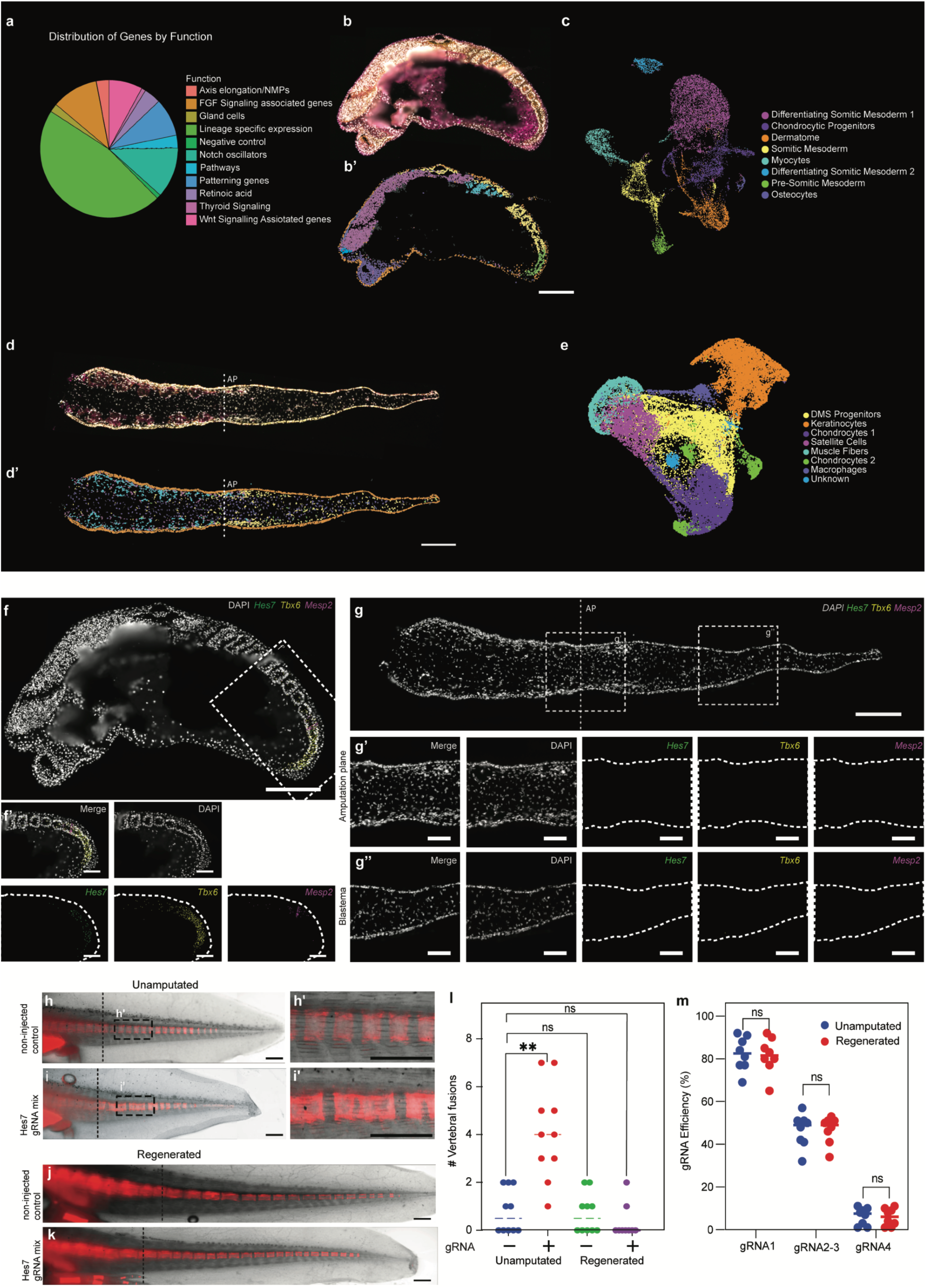
Absence of key somitic clock hallmarks during axolotl tail regeneration. **a,** Distribution of 100 genes selected for Xenium spatial transcriptomics based on their function. **b-c,** Analysis of a representative sagittal section from a stage-28 embryo showing histology (b), spatial distribution (b’), and UMAP distribution (c) of identified cell types. **d-e,** Analysis of a representative coronal section from a 2- weeks post-amputation (wpa) tail blastema showing histology (d), spatial distribution (d’), and UMAP distribution (e) of identified cell types. **f-g,** Xenium expression patterns for *Hes7*, *Tbx6*, and *Mesp2* in stage-28 embryo (f), and 2-wpa tail blastema (g). Dashed line indicates amputation plane (ap). **h-k,** Stereoscopic images of Alizarin red S labeling of segmented vertebrae in uninjured noninjected control (h), Hes7 gRNA mix-injected axolotl (i), regenerated non-injected control (j), and regenerated Hes7 gRNA mix-injected axolotl (k). **l,** Scoring of vertebral fusion effects in the tail (h-k) (n=10). **m,** gRNA efficiency of all four Hes7 gRNAs in unamputated and regenerated tails. Statistical analysis: l: Kruskal-Wallis test, followed by Dunn’s test. m: one-way ANOVA, followed by Šidák correction. ns: not significant, ** p< 0.01. Dashed lines indicate amputation plane (ap). Scale bar: b, d, f-k: 500 μm. f’-g’’: 200 μm.

In axolotl embryos, we observed polarized somitic expression of resegmentation markers *Tbx18* and *Uncx4.1*, but neither was expressed in the tail blastema (Extended Data Fig. 1d-e, 2g, n, u). Hematoxylin and eosin (H&E) staining and the absence of *Tbx18/Uncx4.1* rostro-caudal polarity strongly argue against classic epithelial somites at blastema stages. Together, these results show that while the tail bud and tail blastema exhibit some similarities in their expression of genes associated with *Fgf*, *Wnt*, and RA signaling, we did not detect somite like epithelial units in the blastema via H&E staining.

## Hes7 is dispensable for vertebral segmentation during regeneration

To determine directly whether somitogenesis-dependent patterning takes place during regeneration, we generated a somitic-clock mutant axolotl through targeted mutagenesis of *Hes7*. The Hairy and enhancer of split (Hes/Her) family of bHLH transcription factors is a family of highly conserved oscillatory genes involved in somitogenesis, including *Hes7* (Extended Data Fig. 3a, b), which shows variable expression in the axolotl tail bud consistent with a role as a somitic oscillator (Fig. 2f)^13^. *Hes7* mutant mice display aberrant somite-boundary formation, resulting in vertebral-segmentation defects^14,15^. To generate *Hes7* mutant axolotls, we injected eggs with Cas9/gRNA complexes representing a mixture of four gRNAs targeting all three exons of the *Hes7* gene sequence (Extended Data Fig. 3b). We were unable to recover homozygous *Hes7* mutant axolotls, as they are apparently embryonic lethal. We could, however, analyze mosaic F0 crispants, which escaped embryonic lethality. We imaged vertebrae of F0 crispants using Alizarin Red S in live, 4-5-cm-long animals. As in mice, these mutant axolotls displayed irregular vertebral segmentation and fusions (Fig. 2h, i) consistent with a role for *Hes7* in somitogenesis during embryonic development. Surprisingly these mutant tails recover a wildtype phenotype following amputation, at which time the tails display regular vertebral segmentation and the normal number of tail vertebrae (Fig. 2j-l, Extended Data Fig. 3c-e). To ensure that this observation did not result from positive selection of wildtype and heterozygotic cells in the regenerate, we analyzed editing efficiencies at each exonic target site in the unamputated tail close to the amputation plane and in the regenerated tails. We found no significant difference in editing efficiency among the target sites (Fig. 2m), indicating that, vertebral segmentation during axolotl tail regeneration proceeds without a detectable requirement for *Hes7* under mosaic loss-of-function conditions. In summary, we show that while vertebral segmentation takes place during tail regeneration, it does so through a non-canonical somite-independent program.

## Somite-like lineage potential in the regenerating tail

We next set out to identify the source of the newly regenerated musculo-skeletal tissues in the axolotl tail. To survey the potency of cells that regenerate the tail, we performed clonal barcode lineage tracing in the regenerated axolotl tail using a CellTag-based strategy adapted to a third-generation foamy virus vector system^16–18^. We generated three unique libraries, each containing a 9-nt barcode and a defined 6-nt tag (V1, V2, V3) (Fig. 3a, Extended Data Fig. 4a-c) associated with the 5’ end of a GFP-encoding insert. High-titer viral libraries (Extended Data Fig. 4d) were used to infect each tail four days prior to amputation (V1), as well as four (V2) and seven (V3) dpa (Fig. 3b, c). We recovered V1, V2, and V3 barcodes from single-cell RNA-sequencing (scRNA-seq) data of flow-sorted GFP^+^ cells at 21 dpa (Fig. 3d-e, Extended Data Fig. 4e-h). To identify putative clones, we calculated the Jaccard similarity index of barcodes detected in cells with somite-derivative signatures (tenocytes, intermediate fibroblasts, dermis, fin mesenchyme, chondrocytes, and myogenic cells) because of fate-mapping studies presented in the next section (Fig. 3f). This analysis identified clones ranging in size from 2-6 cells, which suggests an approximately 6-19% recovery rate when compared to an expected average clone size of 32.89 (V1), 16.91 (V2), and 10.27 (V3) cells, assuming a blastema cell-cycle length of 100 hours^9^. Of the 48 clones analyzed, 39 contained diverse combinations of 2-4 cell types (Fig. 3f). Notably, myogenic cells showed a clonal relationship with each of the other five cell types. Moreover, computational mixing of the two replicates and subsequent clone-calling resulted in correct annotation of 47 of 48 clones, indicating a sufficient diversity of barcodes (Extended Data Fig. 4h-i). Although we did not recover any one clone containing all six cell types, these data are compatible with a model in which a multipotent progenitor in the tail that has potential similar to that of pre-somitic mesoderm, and that may give rise to Dermo-Myo-Sclerogenic (DMS) lineages during regeneration (Extended Data Fig. 4j).

**Figure 3:**
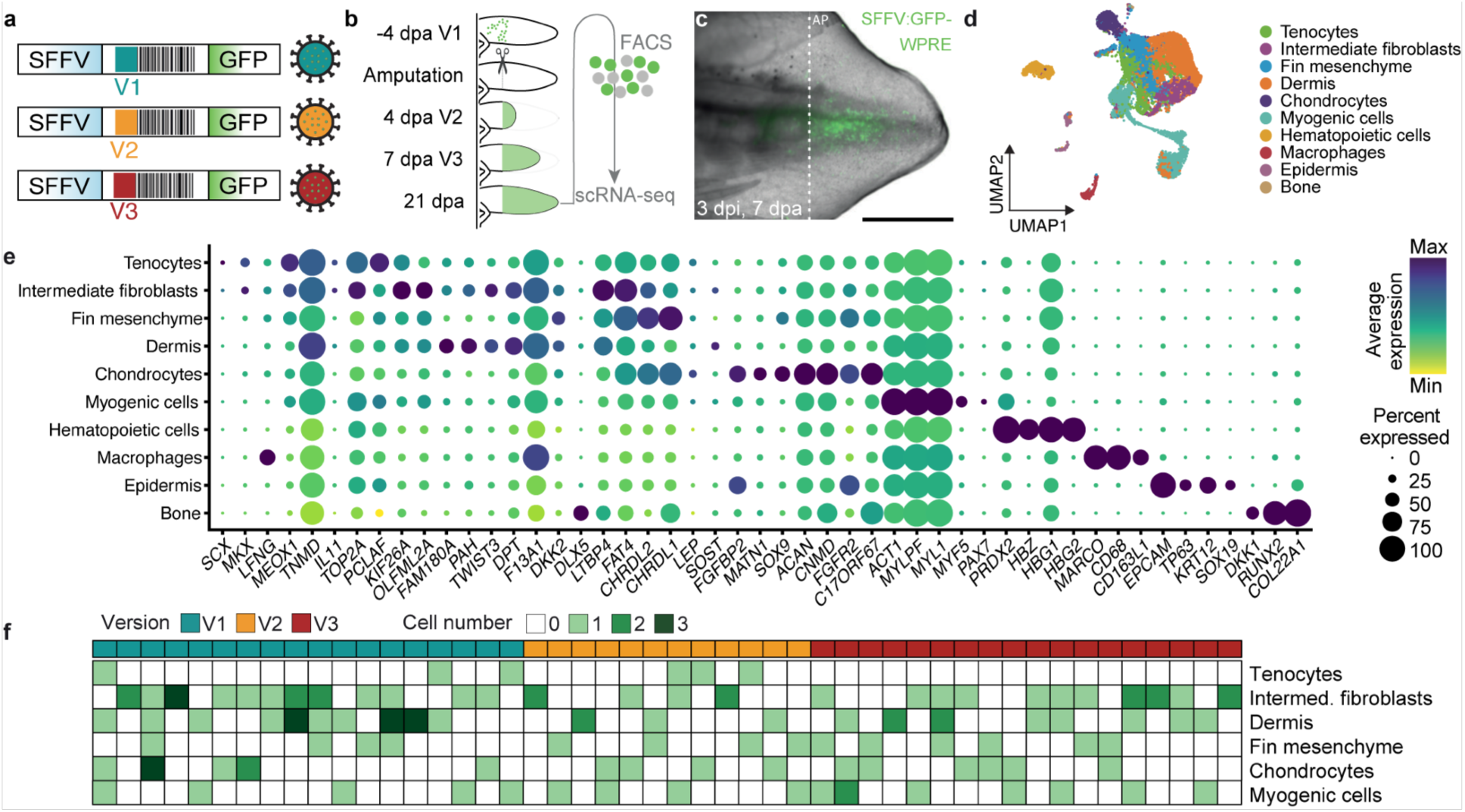
The axolotl tail displays a somite-like lineage potential. **a,** schematic overview of foamy virus barcode labeling vectors. **b,** Experimental overview of infection, amputation and harvesting timepoints. **c,** Representative image of a 7-dpa foamy-virus-infected axolotl tail at 3 days post infection (dpi). **d-e,** UMAP and dotplot representation of different clusters identified in GFP+ cells from foamy-virus-infected tails combining two independent replicates. **f,** Matrix-based depiction of identified clones with a Jaccard similarity index of ≥0.45 (barcodes present in ≥ 2 cells and ≥ 2 UMIs per cell). Each row represents a cell type. Each column represents a clone. Dashed lines indicate amputation plane (ap). Scale bar: c: 500 μm.

## Intermyotomal cells are the source of multipotent musculo-skeletal progenitors

We next aimed to identify and characterize the origin and nature of these progenitors, hypothesizing that they may arise at least in part from connective tissue. We first grafted vertebrae from a constitutively expressing *CAGGs:GFP* transgenic axolotl into an unlabeled host axolotl and observed minimal contribution from vertebrae cells (Extended Data Fig. 5a-d), eliminating vertebrae as a cell source. We then performed Cre/*lox*P genetic fate mapping of tail regeneration using two connective-tissue-specific *Cre-ERT2*-expressing drivers (*Col1A2* and *Twist3*) in combination with a *CAGGs:LoxP-STOP-LoxP-Cherry* reporter, and inducing recombination 10 days prior to tail amputation. Prior to amputation, the *Col1A2* driver broadly labeled the tail connective-tissue population (fin mesenchyme, dermal fibroblasts, tenocytes, and periskeletal and skeletal cells) (Fig. 4a). The *Twist3* driver labeled a smaller subset of tail connective tissue including fin mesenchyme, dermal fibroblasts, and skeletal lineage, but showed very limited labeling of connective-tissue cells residing at the intermyotomal boundaries (Fig. 4b). After tail amputation, the *Twist3* driver showed limited participation of the labeled cells to the regenerate (Fig. 4d, f, Extended Data Fig. 5e-h). In contrast, the *Col1A2* driver contributed labeled cells to regenerated fin mesenchyme, tenocytes, and vertebrae as well as muscle (Fig. 4c, e, Extended Data Fig. 5e-h), consistent with the notion of a common source cell for connective-tissue and muscle lineages. This further excludes dermal fibroblasts and fin mesenchymal cells as the likely cell source and suggests tenocytes as the likely cells source. To assess whether muscle labeling is due to non-specific labeling of the myogenic lineage, such as satellite cells, we performed sectioning and immunostaining with the muscle-progenitor marker PAX7. In converted, unamputated tails we observed no colocalization between Cherry^+^ cells and *PAX7^+^* muscle progenitors (Extended Data Fig. 5i). However, regenerated tails showed the presence of Cherry^+^/PAX7^-^, Cherry^-^/PAX7^+^, and Cherry^+^/PAX7^+^ populations in addition to Cherry^+^ muscle fibers, suggesting that *Col1A2*-labeled Cherry+ cells give rise to at least some PAX7+ muscle progenitors (Extended Data Fig. 5j).

**Figure 4:**
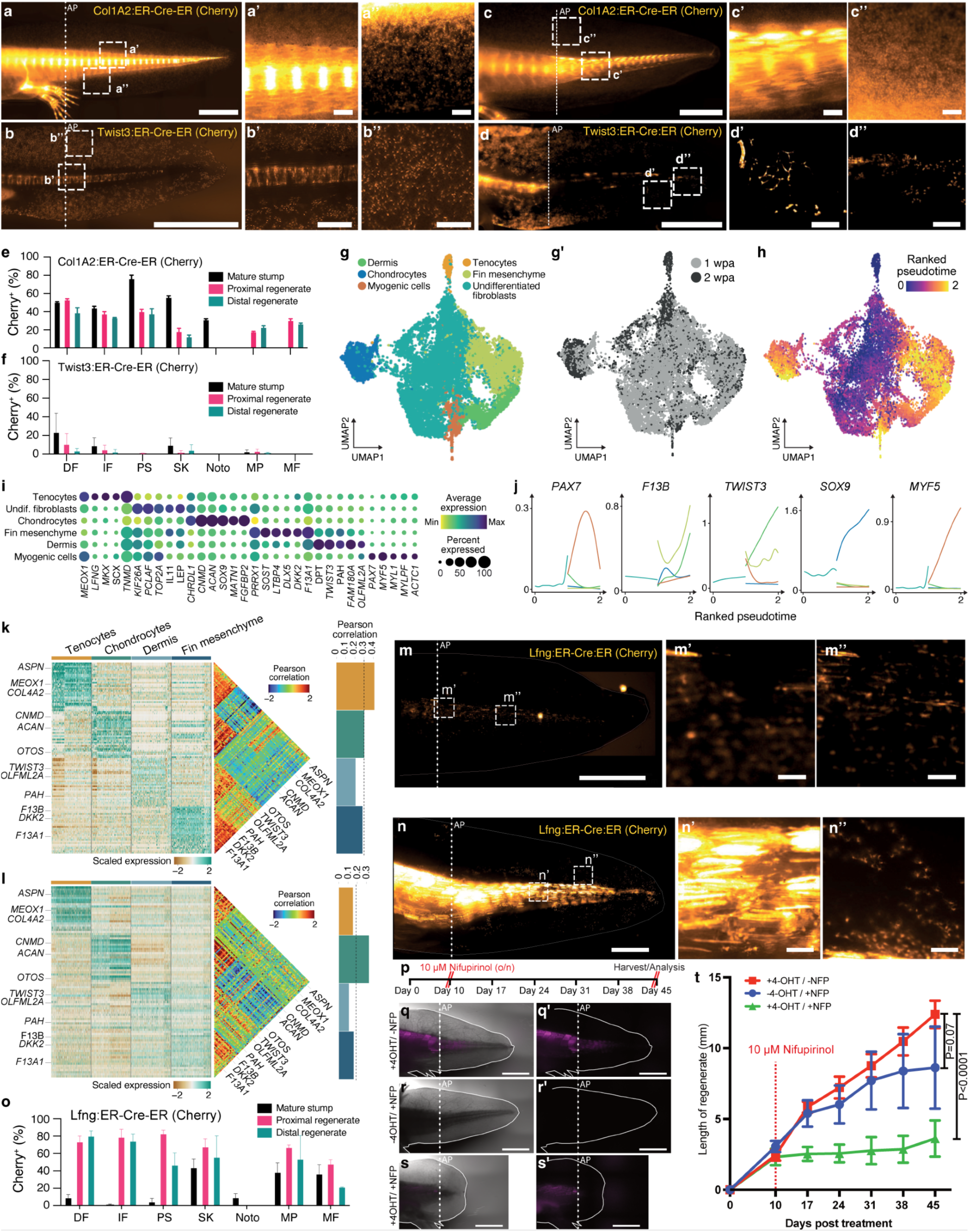
DMS-progenitors are necessary for tail regeneration. **a-d**, Labeling of 4-OHT-treated *Col1a2:ER-Cre-ER;Caggs:lp-Cherry* and *Twist3:ER-CreER;Caggs:lp-Cherry* axolotl tails before amputation (a,b), and at 6 wpa (c,d). **e,f**, Quantification of cell-type distribution at three different positions (mature stump, proximal and distal regenerate, Extended Data Fig. 3g) of the *Col1a2:ER-Cre-ER;Caggs:lp-Cherry* (e) and *Twist3:ER-CreER;Caggs:lp-Cherry* (f) regenerated tails. **g,g’,** UMAP integration of cells from 7 and 14 dpa regenerated tails colored by cell-type identity (left) and by timepoint (right). **h**, Dotplot representation of cell-type marker genes (columns) across the cell type (rows) identified in g. **i**, Pseudotime estimates by a diffusion-map analysis are visualized on the UMAP embedding. **j**, Line plots represent the smoothed gene expression along their pseudotime separated by cell types. **k**,**l**, Heatmap for the top 30 markers (rows) of each cell type (columns) in the uninjured scRNA-seq data (see Extended Data Fig. 6a,b). The genes were used as input for a gene-correlation analysis visualized as a heatmap with average values within each cell type summarized as a barplot (right). Mature tail (k) and 14-dpa blastema (l) **m,n,** Labeling of 4-OHT-treated *Lfng:ER-Cre-ER;Caggs:lp-Cherry* axolotl before amputation (m) and at 6 wpa (n). **o**, Quantification of cell-type distribution of the regenerated *Lfng:ER-CreER;Caggs:lp-Cherry* tail, including the mature stump, proximal and distal regenerates. DF: dermal fibroblasts, IF: interstitial fibroblast, PS: peri-skeleton, SK: skeleton, Noto: notochord, MP: muscle progenitor, MF: muscle fiber. (Extended Data Fig. 3g). **p,** Schematic depiction of nifurpirinol cell-ablation approach. **q-s**, Tail regenerative response after nifurpirinol-based cell ablation in *Lfng:ER-Cre-ER;Caggs:lp-Cherry* transgenic axolotl. 4-OHT treatment (p, p’), and nifurpirinol treatment (q, q’) alone do not affect regeneration; only when both 4-OHT and nifurpirinol treatments are combined is tail regeneration affected (r, r’). **t**, Length of the regenerating tails shown in p-r. Dashed lines indicate amputation plane (AP). Scale bars: a-d, p: 500 μm; a’-d’’, p’: 50 μm. m-n: 1,000 μm. m’-n’’. 100 μm. p-s’: 4,000 μm.

To molecularly pinpoint the potential source cells, we generated scRNAseq data of flow-sorted cells from both the *Col1A2* and *Twist3* driver lines (Extended Data Fig. 6a) and annotated six connective-tissue cell types (tenocytes, fibroblasts, chondrocytes, fin mesenchyme, dermis, and periskeleton) and corroborated cell identities by HCR *in situ* hybridization (Extended Data Fig 6b-h) to localize fin mesenchyme (*Prrx1^+^*), dermal fibroblasts (*Prrx1^low^/Twist3^+^*), periskeletal cells (*Chrdl1^+^*), chondrocytes (*Sox9^+^*), and tenocytes (*Scx^+^*) at intermyotomal boundaries. Consistent with tenocytes as the potential source cells we found tenocytes were depleted in our *Twist3* lineage labelled populations. We found that genes expressed by tenocytes include *Lfng* and *Meox1 (*Extended Data Fig. 6b), which are also expressed during somitic mesoderm development (Extended Data Fig. 1g), in addition to canonical tenocyte markers such as *Tnmd, Scx*, and *Mkx* (Fig. 4i). Immunogold labeling of *Col1a2*-labeled cells identified Cherry+ cells between myomeres that interact with muscle-fiber ends through collagen deposition (Extended Data Fig. 6i-j). Further, whole-body light-sheet imaging of *Col1A2* animals stained for PCNA (proliferative cells) showed that Cherry^+^/PCNA^+^ cells reside in the intermyotomal space in mature tails (Extended Data Fig. 6k). Together, our results suggest that tenocytes labeled by *Scx, Meox,* or *Lfng* are a source population for dermo-, myo-, and sclerogenic lineages respectively in the tail. In addition, based on these results and those of the viral barcoding experiment, we hypothesize that these tenocytes represent multipotent DMS-progenitors in the mature tail.

To assess the trajectories and molecular features by which DMS-progenitors differentiate during tail regeneration, we performed scRNA-seq analysis on the *Col1A2*-labeled lineages at one and two weeks post-amputation (wpa) (Fig. 4g). Consistent with our hypothesis, trajectory analysis implicated tenocytes as DMS-progenitors (Fig. 4 h, j), and later blastema time points showed emergence of the myogenic lineage in the *Col1A2* driver dataset (Fig. 4g’). The transcriptional signatures of all blastema cell clusters were similar to those identified in the mature tail (Fig. 4k, l). This suggests that no unique cell cluster emerges during regeneration, and that DMS-progenitors represent a resident progenitor population of the mature tail. Using HCR-ISH of *Lfng^+^/Meox1^+^* as a marker for DMS-progenitors, we examined their localization during regeneration (Extended Data Fig. 7a-f). At 1 wpa, we found accumulation of DMS-progenitors throughout the blastema (Extended Data Fig. 7a-c), while at 2 wpa, myomeres separated by DMS-progenitors emerged (Extended data Fig. 7d-f).

To test whether *Lfng*-expressing cells in the intermyotomal space indeed represent source cells for regeneration, we generated an *Lfng*-driven Cre-ERT2-expressing transgenic axolotl line. Conversion of the *Lfng* driver 10 days prior to regeneration showed that Cherry expression was absent in the fin mesenchyme and was restricted to the inter-myotome population of the medial axolotl tail (Fig. 4m). Upon tail amputation, the *Lfng* descendants contributed extensively to all somitic mesodermal lineages of the regenerating tail (Fig. 4n, o, Extended Data Fig. 5e-h), suggesting faithful labeling of DMS-progenitors. To examine whether *Lfng-*labeled DMS-progenitors are necessary for tail regeneration, we crossed the *Lfng-*driven CreERT2-expressing transgenic axolotl line to a *lox*P-dependent NTR2.0 (second-generation of nitroreductase) transgenic line (*CAGGS:LoxP-nBFP-LoxP-Cherry-T2A-NTR2.0*^19^). Upon nifurpirinol treatment, we found that ablation of *Lfng*-labeled DMS-progenitors prevented complete tail outgrowth (Fig. 4p-t). We did not quantify ablation efficiency or lineage selectively; therefore, incomplete or off-target ablation effects cannot be fully excluded. Based on these results, together, we conclude that *Lfng^+^* DMS-progenitors contribute to regeneration of somitic lineages and are required for complete tail outgrowth in our ablation paradigm.

## DMS-progenitors represent a non-embryonic identity

Our *Hes7*-mutant and scRNA-seq data suggest that regeneration of somitic mesoderm lineages in axolotl differs from the embryonic program. This would contrast with regeneration of limbs in axolotl, where limb connective-tissue cells dedifferentiate to a state that is molecularly comparable to a limb-bud mesenchyme progenitor^20^. To further investigate this contrast in processes, we integrated all mature and regeneration timepoints into a single UMAP (Extended Data Figure 8a-c) and performed scRNAseq on four tail-bud stages (stage 25, 28, 30, 35). Results showed that, consistent with our earlier Xenium spatial transcriptomic analysis, genes commonly associated with somitogenesis and resegmentation were expressed during embryonic development but did not show increased expression during tail regeneration compared to uninjured tails (Fig. 5a-b). However, we did detect expression of the DMS markers *Lfng* and *Meox1* in the combined mature dataset (Fig. 5b). In addition, we observed expression of two highly conserved markers of resegmentation (*Uncx4.1* and *Tbx18*) in the embryo via scRNAseq, Xenium spatial transcriptomics, and HCR-ISH (Fig. 5b, Extended Data Fig. 1h, 8c, d), but neither technique showed expression of these markers during tail regeneration (Fig. 5b, Extended Data Figure 1i, 2g, n, u 8c, e) Finally, to assess whether tail blastema cells acquire tail-bud-like identities, we performed quadratic programing on published limb^21^ and the above tail development and regeneration datasets. Results show that unlike limb blastema cells which acquire a limb-bud identity, tail blastema cells do not acquire a tail-bud identity (Fig. 5c). In summary, DMS-progenitors have a distinct transcriptional profile and regenerate tail through a process transcriptionally distinct from that used by embryonic somitic mesodermal cells.

**Figure 5:**
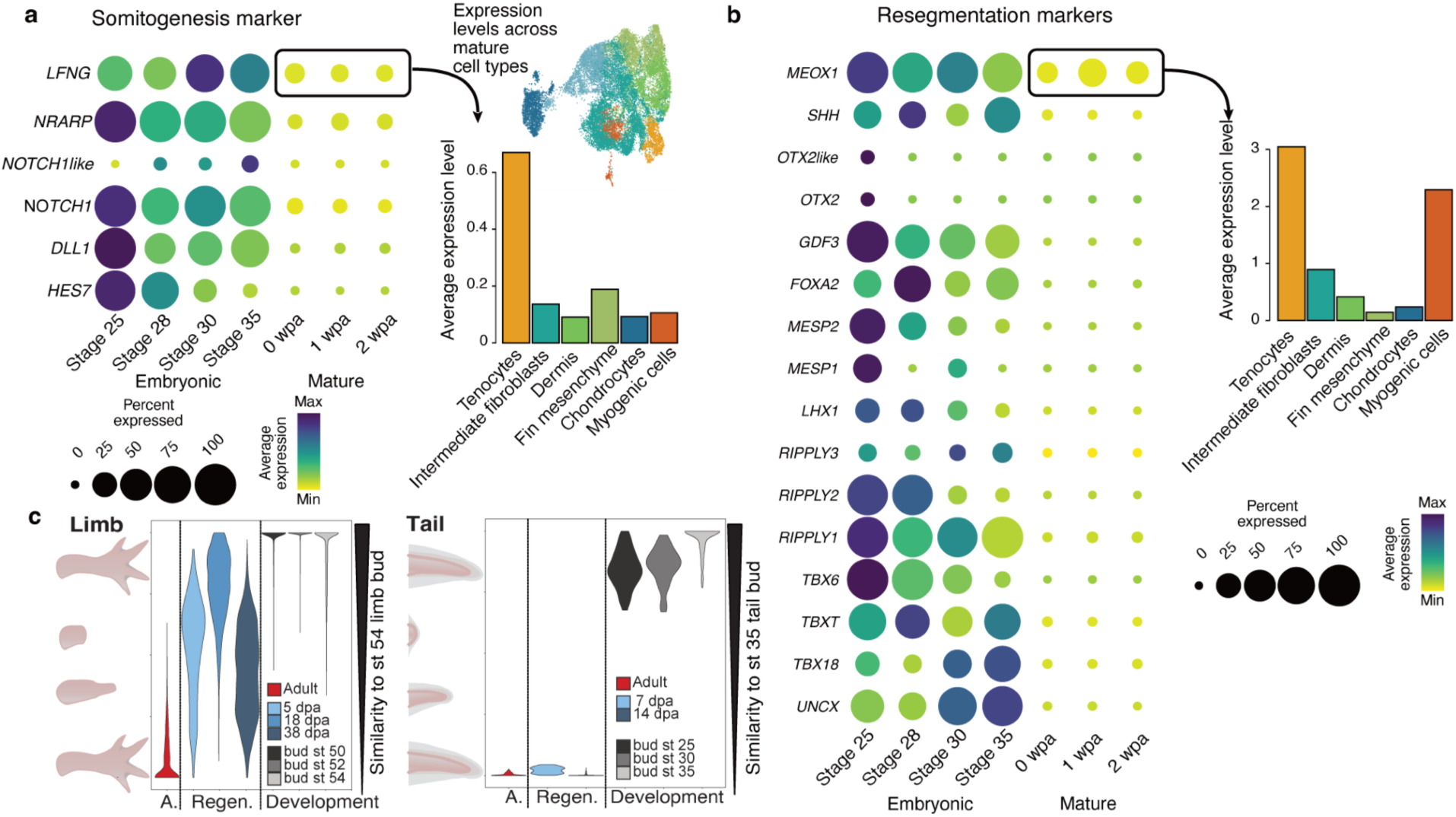
Tail blastema cells do not acquire a tail-bud identity during tail regeneration. **a,b,** Dotplot representation comparing the expression of various somitogenesis (a) and resegmentation (b) markers between tail-bud, mature-tail, and tail-blastema datasets. **c,** Comparative quadratic programming shows that limb cells acquire embryonic limb-mesenchyme identity during limb regeneration (from Lin et al.)^21^ but tail cells do not acquire tail-bud identity.

## Scalable vertebral segmentation takes place via buckling instability

To understand the distinct mechanisms of vertebral segmentation during regeneration, we developed a novel vertebrae extirpation assay, which uncouples vertebral segmentation from axis elongation. By 1 wpi, the injury site collapsed (Fig. 6a, b). Subsequently, by 2 wpi cartilage condensation was observed and the injury site expanded (Fig. 6a, b). By 3 wpi, a stereotypical wrinkling pattern had formed on the outer surface of the regenerating cartilage rod (Extended Data Fig. 9a-g). The wavelength of these wrinkles matches the wavelength of the vertebrae that form at 4 wpi (Fig. 6a, Extended Data Fig. 9a-g) Our combined observations of (i) the collapse of the injury site, and (ii) subsequent expansion of the cartilage rod and wrinkling before vertebrae patterning, suggest a buckling instability due to compression, a mechanism that has been proposed to underlie pattern formation during development of tubular structures in different organisms^22–25^. Buckling instability is a mechanics-based process of periodic pattern formation, which, unlike morphogen- or cell-based processes, is not intrinsically constrained in its scaling. This property is particularly relevant given that axolotl can regenerate their tails at all stages of their life. In this model, under weak compression or a lack of it, a rod maintains a straight and stable shape. Above a critical level of compression, stresses are relaxed locally, producing periodic surface wrinkling, with a wavelength that depends only on a few geometrical and mechanical parameters of the rod. More specifically, we developed a mechanical theory for growth in confined elastic tubes comprising a shell and a core, suggesting two regimes of wrinkling instability (See materials and methods section ‘Buckling instability models’ for details): i) a geometry-dominant model for tubes with very soft cores, which exhibit very little mechanical resistance to compression; in this model, the wavelength (λ, which in our theory represents the distance between vertebrae) depends only on tube geometry, scaling as 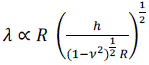 (*λ*: wavelength, *R*:radius, ℎ: shell height, *ν*: Poisson ratio) and not on any stiffness parameters; and ii) a stiffness-dominant model for tubes with cores that have a non-negligible stiffness; in this model, the wavelength is dependent on the relative thickness of the shell and the core, scaling as 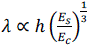 (*λ*: wavelength, ℎ: shell height, *E*_*s*_: shell Young’s modulus, *E*_*c*_:core Young’s modulus), similar to previous theories of buckling induced wrinkling during the formation of crypts and villi, and epithelial folding^25–28^. To determine which model best fits vertebral segmentation during axolotl regeneration, we first measured the mechanical properties of the regenerating tube, i.e., the axolotl cartilage rod, at 4 wpi, using atomic force microscopy (AFM). We found that the mean Young’s modulus of the core is 1.7 kPa, and that of the shell is 6.9 kPa, which together demonstrate a soft core and stiffer shell (Fig. 6c, e, Extended Data Fig. 9h, i). We then quantified the shell thickness (h) and tube radius (R), obtaining mean values of h = 91 μm (Fig. 6f) and R = 450 μm (Extended Data Fig. 9j). Together, these measurements are consistent with an intermediate regime between the two models described above, with a theoretical wavelength of ∼500 μm, closely matching our experimental observations in a parameter-free manner (Fig. 6h, Extended Data Fig. 9g). The geometry-dominant model predicts that the wavelength (*λ*) should increase as the cartilage rod radius (*R*) increases. To test this we took advantage of the variable vertebrae length across the axolotl tail by recording both the wavelength (*λ*) and the diameter for each newly formed vertebra at 3 different amputation planes (4, 10, and 16 myomeres posterior to the cloaca) and used this to plot *λ* and radius 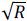. We found that regardless of the amputation plane, our simple geometric prediction of *λ* ∝ 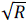 provides an excellent fit for the data (Fig. 6g, h).

**Figure 6:**
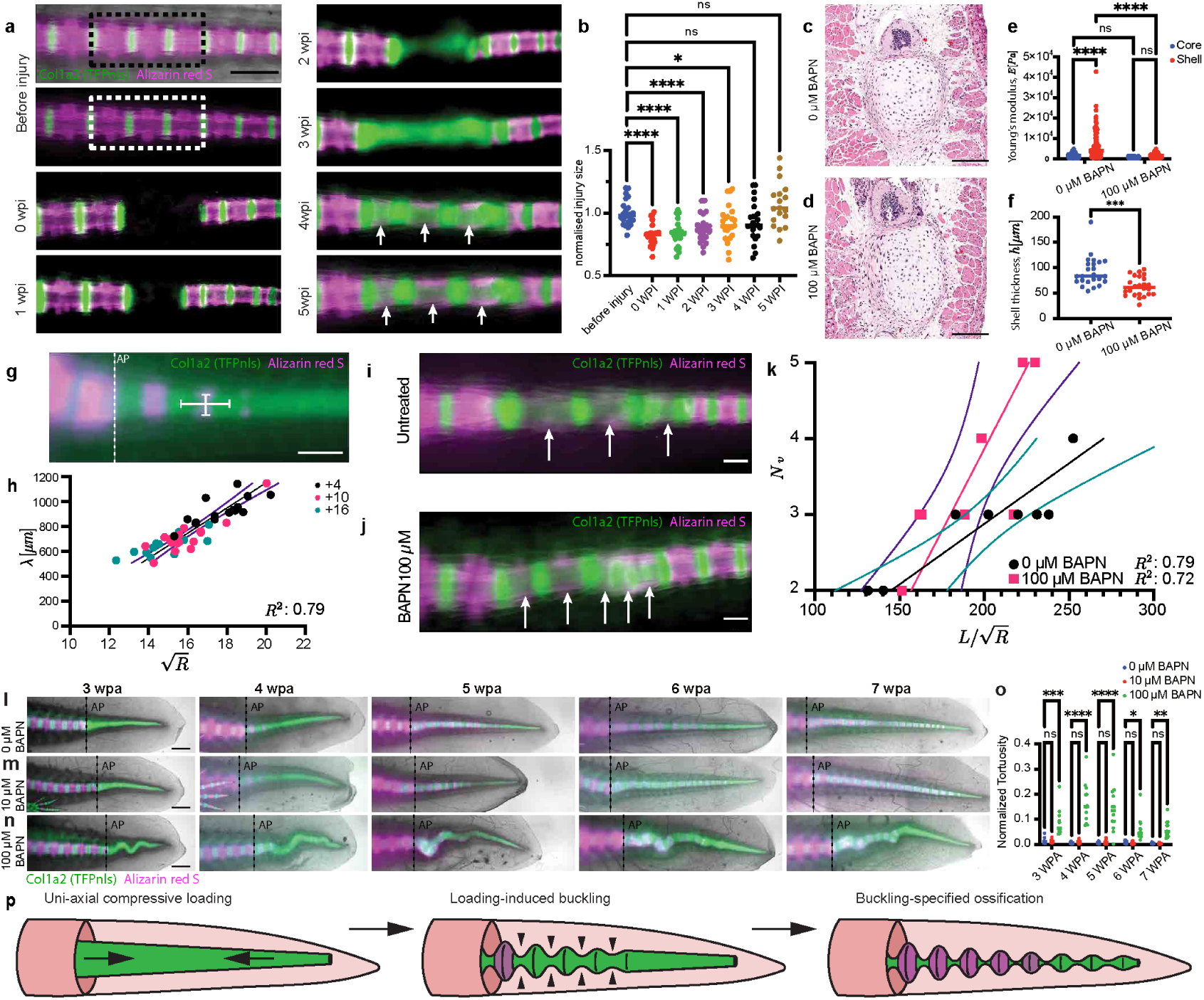
Vertebral segmentation during tail regeneration takes place through buckling instability. **a**, Stereoscopic images of local injury time series in *Col1a2:TFPnls* (green) axolotl labeled with Alizarin red S (magenta). Three-vertebrae injury site is marked using a dashed box. **b**, Quantification of injury-site length over a 5-week period, relative to the average injury-site length at 0 wpi. **c-d**, H&E stain of a 4-wpi cross-section after local injury untreated **(c)** and 100 μM BAPN-treated **(d)**. **e**, Young’s modulus of the core and shell with and without BAPN treatment. **f**, Quantification of shell thickness with and without BAPN treatment. **g**, Stereoscopic image of *Col1a2:TFPnls* (green) axolotl labeled with Alizarin red S (magenta) at 4 wpa, depicting the recording of wavelength and diameter during tail regeneration. **h**, Quantification of *λ* and 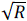 of newly added vertebrae throughout a 7-week regenerative response. Tails were amputated at +4, +10, and +16 myomeres posterior to the cloaca. **i-j,** Regenerated vertebrae at 5 wpi without **(i)** and with **(j)** 100 μM BAPN treatment. **k**, Quantification of the number of vertebrae at 5 wpi in the local injury relative to the aspect ratio of the regenerated cartilage rod (*L*/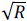) **l-n**, Tail regeneration time series treated with 0 (l), 10 (m), and 100 μM BAPN (n). **o,** Normalised tortuosity of the regenerated cartilage rod in l-n, Normalised tortuosity is calculated 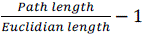. Statistical analysis: b: one-way ANOVA, followed by Šidák correction, e: two-way ANOVA, followed by Šidák correction, f: unpaired t test. h, k: simple linear fit, dashed lines indicate 95% confidence interval, o: two-way ANOVA, with Dunnett’s multiple comparison. Dashed lines indicate amputation plane (ap). Arrows indicate regenerated vertebrae. Scale bar: a, g: 1,000 μm. c-d: 250 μm. i-j: 500 μm. l-n: 2,000 μm.

A key prediction of the stiffness-dominant model is that altering the relative contributions of the shell and core mechanics should change the vertebral wavelength (*λ*). To test this, we treated the regenerating cartilage rod in the vertebrae extirpation assay with β-aminopropionitrile (BAPN), an inhibitor of LOX-mediated collagen crosslinking^29,30^. AFM measurements at 4 wpi demonstrated that the treatment with 100 μM BAPN significantly lowered the shell’s Young’s modulus and reduced the shell thickness (Fig. 6c-f, Extended Data Fig. 9h, i), both predicting a reduced wavelength, which should result in a larger number of vertebrae within a given length unit. Indeed, we found that treatment with BAPN resulted in a larger number of regenerating vertebrae compared to control in the extirpation model (Fig. 6i-k). Furthermore, during tail regeneration, transient treatment with 10 μM BAPN induced a transient reduction in the *λ*/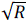 ratio, highlighting the progressive nature of buckling-instability-guided vertebrae patterning during tail regeneration (Fig. 6l, m, Extended Data Fig. 9k). Interestingly our theory assumes that external constraints from surrounding tissues prevent other modes of buckling, such as global deformation of the entire rod, which is expected under free boundary conditions. To test this, we treated the regenerating tail where one end of the rod is free with 100 μM BAPN. Strikingly, this induced global spatial oscillations of the entire rod (Fig. 6l-o) that are fully consistent with our buckling model (Fig. 6j)^31,32^, and demonstrates that the cartilage rod is under a compressive load in both regeneration paradigms.

Together, these results support a quantitatively predictive buckling-instability model for vertebral segmentation during regeneration, a mechanism that inherently confers scalability to the system during regeneration (Fig. 6o).

## Conclusions

Here we show that regeneration of somitic mesodermal lineages in the axolotl tail proceeds through mechanisms that are both morphologically and molecularly distinct from those that take place during embryonic development. In contrast to embryonic somitogenesis, the somitic clock gene *Hes7* plays no detectable role in vertebral segmentation during regeneration. Instead, segmentation emerges through a mechanically driven buckling instability. We identify the cellular source of regenerated somitic mesoderm as *Lfng*⁺/*Meox1*⁺ tenocytes, which we term Dermo-Myo-Sclero (DMS) progenitors, located at the intermyotomal space. This regenerative strategy does not closely mirror axolotl limb regeneration, where fibroblasts broadly dedifferentiate to an embryonic limb-bud–like state^20,21^; rather, the tail regenerative strategy is similar to neural regeneration, where defined adult stem cells undergo lineage-restricted neurogenesis^33^. Previous lineage-tracing studies in larval zebrafish identified a *Col1a2*-expressing connective-tissue cell population capable of contributing to muscle differentiation^34,35^, potentially through activation of MEOX1^36^. However, because adult zebrafish do not regenerate their primary body axis, the relevance of this progenitor-like state to vertebral segmentation and axial regeneration in zebrafish remains unresolved.

Although buckling instability has not been previously proposed as a mechanism for vertebral segmentation, our findings show that this model is a particularly good fit for vertebral segmentation during regeneration. Specifically, because the wavelength of buckling is set by tissue geometry and material properties, vertebral segmentation can automatically scale with the size and growth state of the regenerating tail. This provides a robust means of generating proportionate, periodic structures during large-scale tissue regeneration without reliance on a fixed molecular prepattern. Such mechanically driven scaling mechanisms may represent a general strategy for regenerating other large, periodic anatomical structures and warrants investigation beyond the vertebral column.

Our findings show conceptual parallels with vertebral patterning during zebrafish development^37–40^. In crown teleosts, including zebrafish, classical *Tbx18/Uncx4.1*-based resegmentation is lost, and vertebral centers are specified through a mechanically mediated process during which vertical myosepta transmit tension to the notochord, leading to periodic induction of vertebral elements. While axolotl tail regeneration and zebrafish development employ distinct molecular players, both rely on mechanical forces that reshape the tissue precursor from which the vertebral column emerges. Whether this similarity reflects convergent evolution or a shared ancestral mechanism remains unclear. Tail regeneration is widespread across Euteleostomes, spanning both Sarcopterygii and Actinopterygii. Euteleostome species capable of tail regeneration include the banded knifefish, African lungfish, the extinct early tetrapod *Microbrachis pelikani,* and lizards^41–45^. Notably, vertebral segmentation has been documented in African lungfish and in *Microbrachis pelikani*, but has not been observed in lizards^41,42,44,45^.

Conceptually, our work reveals that segmented vertebrae can arise through two fundamentally distinct pathways: an embryonic, clock-based mechanism and a post-embryonic regenerative mechanism. This distinction has important implications for regenerative medicine, as it suggests that faithful reconstruction of complex axial structures may not require reactivation of embryonic programs. Instead, exploiting mechanically guided patterning of adult progenitor populations may offer an alternative and potentially more accessible route to vertebral column regeneration in other species, including humans.

## Acknowledgements

Acknowledgments

We would like to acknowledge the valuable services provided by the animal facility, Molecular Biology Services, and BioOptics facilities at the IMP and IMBA; Histology, and NGS sequencing core facilities at the Vienna Biocenter; the Comparative Animal Models Core, Light Microscopy Facility (LMF; RRID:SCR_019166), and Comparative Genomics and Data Science Core at MDIBL; MDIBL receives support from the COBRE (P30GM154610) and Maine INBRE grant (P20GM103423) from NIH-NIGMS. Xenium spatial transcriptomics was carried out in the Genomics and Molecular Biology Shared Resource (RRID:SCR_021293) at Dartmouth Cancer Center, which is supported by NCI Cancer Center Support Grant 5P30CA023108 and NIH S10 (1S10OD030242) awards. We thank Dr. Stephen Sampson for proofreading this manuscript.

## Author contributions

Conceptualization: WM, TG, FF, EH, EMT, PM

Methodology: WM, TG, VSJ, FF, RG, RPS, MP, YST, TYL, TK, JW, JFF, FWK, JHG, EH, PM, MR, OGA, PJT, TD, GD, AA, YTS, TYL, TK, JW, EH, BT, DA

Investigation: WM, TG, VSJ, FF, RG, RPS, SCP, MP, JFF, TD, YST, TYL, TK, FWK, JHG, EH, PM, MR, OGA, TD, GD, AA, YTS, TYL, TK, JW, EH

## Materials and Methods

### Animal husbandry, transgenesis, and 4-OH tamoxifen treatment

Axolotls (*Ambystoma mexicanum*) were bred in IMP and MDIBL facilities. All animal handling and surgical procedures were carried out in accordance with local ethics-committee guidelines. Animal experiments were performed as approved by the Magistrate of Vienna and the IACUC committee of MDIBL. “White” refers to a non-transgenic *d/d* strain of axolotl that has leucistic skin due to the absence of melanocytes. Animals were anaesthetized in 0.03% benzocaine (Sigma) before amputation and surgery. Axolotl husbandry, transgenesis, and 4-hydroxytamoxifen (4-OHT) treatment were done as previously described^43^. tgSceI(Mmu*.Col1a2:TFPnls-T2a-ER^T2^-Cre-ER^T2^*)^Etnka^, tgSceI(*CAGGs:LoxP-GFP-LoxP-Cherry*)^Etnka^, tgSceI(*Caggs:LoxP-STOP-LoxP-Cherry*)^Etnka^ and tgSceI(*Caggs:Lp-BFPnls-LP-Cherry-T2a-NTR2.0)^Pmx^* transgenic lines are described previously^20,46,47^. To generate *Lfng* transgenic animals, a mouse *Lfng*^48^ driver element was cloned at the 5′ end of the GFPnls-T2A-ER^T2^-Cre-ER^T2^ (ER-Cre-ER) cassette with flanking ISceI sites. The *Twist3* transgenic animal was generated by knock-in methodology described previously^49^. *Twist3* is a single-exon gene. First, to generate a targeting construct, genomic sequence containing 450 bp upstream seq of the 5’UTR of *Twist3* along with its 5’UTR and ORF (without stop codon) (total 1,230 bp) was amplified from axolotl genomic DNA and was cloned at the 5’ end of the MCS-T2a-GFPnls-T2a-ER^T2^-Cre-ER^T2^ base vector to generate 450bp_upstream_seq-5’UTR-Twist3ORF (without stop codon)-T2a-GFPnls-T2a-ER^T2^-Cre-ER^T2^ construct. Next, embryos were co-injected with the targeting construct, CAS9 protein and gRNA (GCTGCAGCCCATGCAATC-TGG) with a target site 295 bp upstream of the ORF to create dsDNA breaks in both the genome and the targeting construct. Embryos were screened for nuclear GFP fluorescence at the 3-cm stage and grown up as potential breeders. Successful germ-line transmission was evaluated by nuclear GFP fluorescence in dermal fibroblasts across the body, and a line was established. Constructs used for generating transgenic axolotl driver lines are available at Addgene (200104, 200105, 200106, and 200107).

### Xenium spatial transcriptomics

Axolotl embryos were staged to developmental stage 28 after fertilization and fixed overnight in 4% paraformaldehyde (PFA; Sigma-Aldrich, 441244-3KG) at 4°C. Embryos were washed with phosphate-buffered saline (PBS) and incubated in 2% gelatin (Sigma-Aldrich, G2500-100G) at 37°C in a water bath for 1 hour and placed under vacuum for 30 minutes at room temperature. The embryos were washed with ice-cold PBS containing 2% gelatin, and subjected to a sucrose infiltration series (7.5%, 15%, and 30%). For cryopreservation and dehydration, the embryos were equilibrated in a 30% sucrose: OCT compound solution (1:1) and embedded in OCT. Samples were sectioned at a thickness of 20 µm using a cryo-microtome (Leica CM1860 UV). Tails were collected from animals approximately 3 cm in length that had undergone tail amputation and were allowed to regenerate for 14 days. On day 14, the animals were anesthetized with 0.03% benzocaine, and the regenerated tails were harvested using a scalpel (Aspen Surgical Products 371621). The collected tails were arranged and aligned in a cryomold containing OCT compound (Leica Biosystems 14020108926). Once properly positioned, the OCT was rapidly frozen by placing the cryomold on a pre-chilled heat block submerged in liquid nitrogen. Cryosections were prepared at a thickness of 20 µm and subsequently transferred onto Xenium slides for downstream processing. Xenium panel design, tissue histology, and processing were carried out according to the manufacturer’s instructions for preparing Xenium slides from fresh frozen tissues (10x Genomics, Protocol: CG000579), with the following modification: slides containing embryo samples were fixed in methanol for 2hrs at room temperature (rather than the 30min stated in the protocol). Samples were subjected to probe hybridization, ligation, and amplification steps, and stained with the Cell Segmentation Add-On Kit (10x Genomics Protocol: CG000749). Xenium slides were subsequently processed on a Xenium Analyzer instrument running Xenium Software (v.2.0.1.0) and Onboard Analysis Software (v.2.0.0.10), generating the output data bundle including principal component analysis (PCA), K-means and graph-based clustering, differential gene expression analysis, and UMAP (Uniform Manifold Approximation and Projection)-based dimensionality reduction for downstream analyses. A comprehensive list of all detected transcripts is available in GEO: GSE313338. The custom panel was designed to target 100 genes associated with somitogenesis and cell-type classification within the blastema. Each target gene was assigned 6–8 probes, designed based on the provided gene sequences. Final gene selection and probe numbers were validated by cross-referencing with unfiltered single-cell transcriptomic data to ensure balanced representation across cell types and to prevent oversaturation of any single population, which could otherwise obscure distinct fluorescent signals. Slides were stained with hematoxylin and eosin (H&E) using a Sakura Tissue-Tek Prisma stainer and imaged at 40× magnification with an Aperio GT450 scanner (Leica). Images were exported from the Xenium Browser in PNG format, corresponding to subcellular, near single-molecule RNA resolution. Exported images were imported into Fiji (ImageJ, NIH). Each image was converted to 8-bit grayscale and inverted to obtain a dark background, after which the images were binarized. Binary objects were dilated to increase dot visibility.

Xenium cell-feature matrices were processed using a custom Seurat script (version 5.3.0) to generate a combined cluster profile for the blastema and embryonic timepoints. Data were then normalized and scaled with the SC Transform package (version 0.4.2), followed by principal component analysis (PCA) using all genes, with PC cutoffs of 11 and 16. The processed matrices were subsequently analyzed with Find Neighbors and FindClusters, and dimensionality reduction was performed with t-SNE and UMAP for visualization.

Xenium data were manually processed within the Xenium Explorer 3 (version 3.2.0). Sub-epithelial regions of the 14-dpa blastema and embryonic tails were isolated through selections that were restricted to include the blastema/pre-somitic mesoderm and a segment of uninjured/mature muscle. Transcripts and cells collected in these regions were parsed using an in-house algorithm developed using Python3 (version 3.10.14). Within the selection area, cells identified by Xenium Ranger (version 3.1.1) were used to define sample boundaries. Boundary coordinates were analyzed to orient local anatomical features, including the distal and proximal edges of the tail. Xenium data are available at GSE313338.

### Generation of *Hes7* mutants

*Hes7* mutant animals were generated by co-injecting freshly fertilized *d/d* eggs with RNP complexes containing a cocktail of four different guide RNAs targeting all three exons of *Hes7* using standard methods^50,51^. gRNA was prepared using primer templates and *in vitro* RNA synthesis. gRNA efficiency was assessed by targeted amplification of gRNA target sites compatible with Interference of CRISPR Edits (ICE) analysis. In brief, gDNA was prepared by incubating 1-2 mm of tail tissue with 200 μL of 50mM NaOH at 95°C for 15-20 minutes, and subsequently neutralizing the solution by mixing it with 50 μL of 1M TRIS-in 50 mM NaOH. Samples were centrifuged at 15,000 RCF for 6 minutes, and the supernatant was subsequently used for PCR. 1 μL of gDNA was used to seed 50-μL PCR reactions and cleaned up for Sanger sequencing using magnetic beads. Sanger sequencing was performed using a primer that is internal to the amplicon. gRNA efficiency was determined through ICE analysis software (Synthego, Redwood City). The following primers were used

sgRNA generation:

Common_sgRNA_reverse: 5’AAAAGCACCGACTCGGTGCCACTTTTTCAAGTTGATAACGGACTAGCCTTATTTTA ACTTGCTATTTCTAGCTCTAAAAC 3’

Hes7_gRNA1: 5’GAAATTAATACGACTCACTATAGGGTCGGTCAGCTTCATGGTGTTTTAGAGCTAGAA ATAGC 3’

Hes7_gRNA2: 5’GAAATTAATACGACTCACTATAGGGCATGAACCACAGCCTGGGTTTTAGAGCTAG AAATAGC 3’

Hes7_gRNA3: 5’GAAATTAATACGACTCACTATAGGTCAGTGGCTTCCAGAAGCGTTTTAGAGCTAG AAATAGC 3’

Hes7_gRNA4: 5’GAAATTAATACGACTCACTATAGGGGGGTTCCGGAATTTCTGGTTTTAGAGCTAGA AATAGC 3’

Sequencing:

Hes7_gRNA1_Fwd:5’ AACTAGCTCAAACCGGCAGA 3’

Hes7_gRNA1_Rev:5’ GAGGCCCGAACAGATATTGA 3’

Hes7_gRNA1_Seq:5’ CTCCTGCTGGGGAAGTCAT 3’

Hes7_gRNA2+3_Fwd:5’ TGATGCTGTTCCCGTGATAA 3’

Hes7_gRNA2+3_Rev:5’ AGGAGGTTGGACCTCTTGGT 3’

Hes7_gRNA2+3_Seq:5’ CACGTTGCAGGATCAGGAAT 3’

Hes7_gRNA4_Fwd:5’ TTTCTCGCCTCACAAGGTCT 3’

Hes7_gRNA4_Rev:5’ AAGTTCCGTTGAGGGTGATG 3’

Hes7_gRNA4_Seq:5’ GTGGTGAGGCTGATCCATCT 3’

### BAPN treatment

3-Aminopropionitril fumarate (Sigma-Aldrich, A3134-25G) was dissolved in tap water to generate a stock solution of 100 mM, aliquoted and stored at −80 °C. Aliquots were further diluted into tap water to reach working solutions of 10 and 100 μM. Animals were bathed in BAPN solutions and were refreshed every other day.

### Cell ablation using NTR2.0

Transgenic axolotls were generated by crossing *Lfng:GFPnls-T2A-ER^T2^-Cre-ER^T2^* and *CAGGs:loxP-BFPnls-loxP-Cherry-T2a-NTR2.0* lines to enable tamoxifen-inducible NTR2.0 expression in *Lfng-*positive cells. Animals (∼3.5 cm, 3 months old) were housed in Holtfreter’s solution at 19°C under a 12-hour light/dark cycle. NTR2 expression was induced by treating animals with 2 µM 4-hydroxytamoxifen (4-OHT, Sigma# H6278-50MG) via 12-hour immersion on days 0, 2, and 4, followed by maintenance in 20% Holtfreter’s until day 10, designated as 0 days post-conversion (dpc). At 0 dpc, tail amputations were performed at the fifth myotome posterior to the cloaca under 0.03% benzocaine anesthesia. Animals were pre-assigned into three groups: Group 1 (Nifurpirinol+, 4-OHT–, n=5) as non-converted drug-only controls; Group 2 (Nifurpirinol–, 4-OHT+, n=6) as converted but no-drug controls; and Group 3 (Nifurpirinol+, 4-OHT+, n=7) as the ablation group targeting *Lfng*-lineage cells via NTR2-mediated cytotoxicity. 10 µM Nifurpirinol (MedChem Express, HY-135470) was delivered by 24-hour immersion and prepared from a DMSO stock solution. Imaging was conducted under consistent conditions at 0, 10, 17, 24, 31, 38, and 45 dpc using a Zeiss AxioZoom V16 microscope at 20x magnification following anesthesia in 0.03% benzocaine.

### Histology

Embryos were fixed in Carnoy’s solution, and blastema and mature tissues were fixed in 4% PFA. H&E staining was done using an Epredia Gemini AS Automated Slide Stainer (Fisher Scientific, Waltham) with standard protocols. Stained slides were imaged using a Panoramic 250 FLASH II digital scanner (3DHistech Ltd., New Jersey).

### Lineage tracing and analysis

To perform Cre-*Lox*P lineage tracing experiments, Cre driver lines were crossed with either tgSceI(*CAGGs:LoxP-GFP-LoxP-Cherry*)^Etnka^ or tgSceI(*CAGGs:LoxP-STOP-LoxP-Cherry*)^Etnka^. Single- and double-transgenic animals were grown up to 4 cm snout-to-tail size. Animals were treated three times at alternate days with 2 μM 4-OHT by bathing to obtain indelible Cherry labeling of cells. Labeled animals were amputated 10 days post-conversion (from the first 4-OHT treatment), near the 6^th^ myotome from the cloaca, and contributions of *Cherry^+^* cells to the regenerate were accessed by stereoscopic imaging of a regenerated tail every week for the next 6 weeks using an Axio Zoom.V16 (Zeiss, Jena) equipped with an Orca-Flash4.0 camera (Hamamatsu Photonics, Hamamatsu) and an X-Cite Xylis LED illuminator (Excelitas Technologies, Waltham). At the end of 6 weeks, tails were harvested near cloaca, and IHC or HCR analysis was performed.

### Antibodies

The primary antibodies used in these studies are rabbit anti-RFP (Rockland# 600-401-379, 1:100), mouse anti-COL1A2 (DSHB # Sp1.D8, 1:100), mouse anti-PCNA (Calbiochem#NA03, 1:500), mouse anti-PAX7 (DSHB#Pax7, 1:100), mouse anti-MHC (monoclonal antibody 4A1025, a kind gift from S. Hughes), goat anti-SOX9 (R&D#AF3075, 1:100), rabbit anti-SOX9 (Chemicon#AB5535, 1:1000), rabbit anti-PRRX1^20^ (1:200), and rabbit anti-Laminin (Merck, #L9393 1:200). Secondary antibodies used in these studies were procured from Thermo Fisher Scientific (Waltham).

### Immunohistochemistry and microscopy

IHC staining was done as previously described^20^. Briefly, 10-μm sections were permeabilized using phosphate-buffered saline (PBS) containing 0.3% Tween-20 and blocked with PBS with 0.3% Triton-X100 and 2% normal hose serum (Vector Laboratories # S-2000). Slides were incubated O/N with primary antibodies in blocking buffer in a humidified chamber at room temperature and subsequently incubated with secondary antibodies and Hoechst at a concentration of 0.5 μg/mL. Mosaic images of sections were acquired on an Axio Imager.Z2 (Zeiss, Jena) equipped with an Orca-Flash4.0 camera (Hamamatsu Photonics, Hamamatsu) using a 20x/0.8 plan-apochromat objective. Alcian blue / Alizarin red S staining was performed as previously described^52^. Stained axolotls were photographed using a Nikon D40 DSLR camera with an AF-S DX NIKKOR 18-55mm f/3.5-5.6G VR II objective (Nikon, Tokyo) and an iPad Air (4^th^ gen) (Apple, Cupertino) as a backlight illuminator.

### Identification of different cell types based on histology

Collagen 1 alpha 2 (COL1A2) is an extracellular matrix (ECM) protein, and in axolotl it is expressed in dermal fibroblasts and skeleton cells (28). Staining tail sections with COL1A2 antibodies showed broad labeling of the basal lamina (which demarcates dermal fibroblasts and basal keratinocytes), skeletal and periskeletal structures, as well as fin mesenchyme. Thus, based on co-staining of PRRX1 and COL1A2 along with laminin and Pax7 of sister sections, we identified the following seven distinct connective-tissue subtypes (Extended Data Fig. 4): 1) “Dermal fibroblasts (DF)” - cells underneath the basal lamina that are COL1A2^+^ and PRRX1med; 2) “Interstitial fibroblasts (IF)” - cells scattered across the fin mesenchyme that are COL1A2^+^ and PRRX1^high^ or PRRX1^med^; 3) “Periskeletal cells (PS)” - cells at the periphery of skeleton that are COL1A2^+^ and PRRX1^high^ or PRRX1^med^; 4) “Skeletal cells (Sk)” - cells that reside inside cartilaginous structures and are COL1A2^+^ and PRRX1^low^; 5) “Notochordal cells (Noto)” - large vacuole-containing cells within the embryonic notochord; 6) “Muscle progenitors (MP)” - PAX7^+^ cells scattered between muscle fibers; and 7) “Muscle fibers (MF)” - MHC^+^ or laminin^+^ cells within the myotome region.

### Whole-mount light-sheet imaging of PCNA-labeled cells

In brief, axolotl samples were cleared and labeled using the Deep-Clear protocol followed by imaging on a custom-built light-sheet system^53^. Samples were fixed in 4 % PFA at 4°C overnight. Specimens were washed several times with PBS at room temperature to remove fixative. After PBS washes, specimens were incubated in Solution-1 at 37°C for 1 day under gentle shaking. After the Solution-1 treatment step of tissue clearing and five short PBS washes at room temperature, samples were treated with 10 % sheep serum at room temperature for several hours. Samples were then incubated with primary antibodies in 5 % sheep serum at 4°C (gentle shaking) followed by additional wash steps followed by incubation with secondary antibodies in 5 % sheep serum at 4°C (gentle shaking). Solution-1 consists of 8% THEED (Sigma-Aldrich, 87600-100ML), 5% Triton® X 100 (Roth, 3051.2), 25% urea (Roth, X999.2) and 5% CHAPS (Hopax) in dH_2_O. Solution-2 was prepared by mixing 50% meglumine diatrizoate (Sigma-Aldrich M5266) in PBS (pH 8.5), and the refractive index was adjusted to 1.45.

Images were acquired using a custom-built light-sheet system equipped with an Olympus Plan Achromat 1.0/0.25 objective (N1564200). Sample were illuminated with LightHUB 4 system (Omicron, Germany) containing the following single-mode lasers: 120mW 405nm LuxX laser, 150mW 488nm LuxX laser, 150mW 561nm Cobalt laser, 100mW 594nm Cobalt laser, and 140mW 647nm LuxX laser. Images were recorded in 16-bit quality as Tiff, with a Kinetix camera (Teledyne Photometrics, USA) 3200x3200px; pixel size 6.5 μm. Images were processed with AMIRA (Thermo Scientific, USA) rendering software.

### Immunogold labeling and electron microscopy

Samples were fixed in freshly prepared 4% PFA. Longitudinal 50 mm thick vibratome sections were prepared on a Leica VT 1200 vibratome, and sections with target cells were selected based on their Cherry fluorescence. Vibratome sections were subjected to pre-embedding immunogold labeling using Nanogold and silver enhancement^54–56^. In brief, samples were blocked and permeabilized in 20% normal goat serum (NGS) / PBS / 0.1% Saponin for 2 hours at room temperature followed by incubation in primary antibody (rabbit anti-RFP, Rockland, # 600-401- 379, 1:100) in 20% NGS/0.05% saponin for 2 days at room temperature. After washes in 20% NGS/0.05% saponin, the samples were incubated with goat-anti-rabbit nanogold (Fab-fragments, Nanoprobes, 1:50) in 20% NGS/0.05% saponin overnight at room temperature. After final washes in 20% NGS/0.05% saponin, the samples were postfixed in 1% glutaraldehyde in PBS, washed in water, and silver-enhanced using the SE-Kit (Aurion, 1 hour incubation time), followed by washes in water, post-fixation in 1% osmium tetraoxide/water (1 hour on ice), washes in water, *en bloc* contrasting with 1% uranyl acetate (1h, on ice), washes in water, and dehydration in a graded series of ethanol (30%, 50%, 70%, 90%, 96% ethanol/water, 3x 100% ethanol on a molecular sieve). The samples were infiltrated in mixtures of the Epon substitute EMbed 812 with ethanol (1+2, 1+1, 2+1, 2x pure Epon), flat-embedded on the surface of an empty Epon dummy block, and cured overnight at 65°C. The region of interest (ROI) was identified on semithin sections (1 mm) stained with 1% toluidine blue/0.5% borax, and the ROI in the block was trimmed for ultrathin sectioning. 70-nm sections were prepared using a diamond knife (Diatome) and the Leica UC6 ultramicrotome (Leica Microsystems, Wetzlar). The sections were mounted on Formvar-coated slot grids and contrasted with 4% uranyl acetate/water for TEM. Finally, sections were imaged with a Jeol JEM1400Plus transmission electron microscope running at 80kV acceleration voltage and equipped with a Ruby digital camera (Jeol, Tokyo).

### HCR-ISH

Samples were fixed in freshly prepared 4% PFA, prepared for cryo-sectioning, and sectioned at 80-μm thickness. HCR staining was performed according to the previously published protocol^57^. The probes were designed using probe generator software from the Monaghan lab (https://probegenerator.herokuapp.com/ - the lab is no longer functional) or by the MDIBL Comparative Genomics and Data Science Core team, and probes were ordered from IDT (Coralville) as oligo pools. All other HCR reagents were ordered from Molecular Instruments (Los Angeles).

### Whole-mount HCR-FISH imaging of axolotl embryos

Stage-30 embryos were fixed in freshly prepared 4% PFA for 1 hour at RT. Embryos were washed with PBS for 3 × 5 minutes and depigmented using 3% H_2_O_2_ and 0.8% KOH in DEPC-H_2_O for 20 minutes. Next, embryos were incubated in Solution 1 of the Deep-Clear method^53^ for 1 hr at 37°C with shaking at 500 rpm on a tabletop shaker. Embryos were washed with 5x SSC-0.1% Tween-20 (SSCT) for 3 × 15 minutes at 500 rpm. Probe hybridization and amplification were done according to the manufacturer’s protocol (Molecular Instruments, Los Angeles). Embryos were mounted in Solution-2 of the Deep-Clear protocol and incubated for 1 hr at room temperature to match the refractive index before imaging. Images were acquired using a Spinning-disk confocal unit (CSU-W1, 25 μm pinhole diameter, Yokogawa, Japan) on a Nikon inverted Ti-Eclipse microscope stand (Nikon Instruments Inc., Japan), equipped with a Nikon CFI Plan Apochromat Lambda D 10X/0.45 for whole-embryo imaging and a Nikon CFI Plan Apochromat Lambda 20X/0.75 lens for the cropped images. Alexa 647, Cherry and DAPI fluorescence were excited with 640-nm, 561-nm, and 405-nm laser lines and collected using a 405/488/561/640 dichroic beamsplitter (Yokogawa) with a ET705/72 (Chroma), ET605/52 (Chroma), and ET436/20 (Chroma) emission filter, respectively. Images were acquired at 2048X2048 pixels in 16 bit with a Scientific CMOS <u>Zyla 4.2</u> (Andor Technology, United Kingdom) controlled with NIS AR 5.41 (Nikon Instruments Inc., Japan) software and saved in Nd2 or Tiff file format. A maximum intensity projection (MIP) of the z-stack was created using Fiji software. Images were acquired using a Nikon Ti-E Yokogawa CSU-W1 / Nikon C2+ confocal microscope with a 10X Nikon CFI Plan Apochromat Lambda (NA-0.45) lens for whole-embryo imaging and with a 20X Nikon CFI Plan Apochromat Lambda (NA-0.75) lens for the cropped images. A MIP of the z-stack was created using Fiji-ImageJ software.

### Vertebrae transplantation

To visualize vertebrae, 5-cm *d/d* and *CAGGs:eGFP* animals were soaked in 0.05% Alizarin red S (ARS) for 30-60 minutes and rinsed several times in fresh tap water^58^. Subsequently, axolotls were anaesthetized in 0.03% benzocaine (Sigma) and prepared for surgery. Using a SZX16 stereo-microscope (Olympus, Tokyo) equipped with a Zyla sCMOS camera (Andor, Belfast) and an X-Cite Xylis LED illuminator (Excelitas Technologies, Waltham, MA) fluorescent microscope to visualize vertebrae, and fine forceps, 3-4 vertebrae were carefully extirpated from the *d/d* host. Subsequently, 3-4 vertebrae were dissected from the donor and grafted into the previously prepared host. Axolotls were left to recover for 2 weeks, after which tails were amputated through the GFP^+^ donor vertebrae and lineage-traced for a total of 6 weeks.

### Local injuries

Local injuries were performed on 4-7-cm-long *Col1a2:TFPnls-T2a-ER^T2^-Cre-ER^T2^*animals soaked in 0.05% ARS for 30-60 minutes and rinsed several times in fresh tap water^58^. Subsequently, axolotls were anaesthetized in 0.03% benzocaine (Sigma) and prepared for surgery. Using a SZX16 stereo-microscope (Olympus, Tokyo) equipped with a Zyla sCMOS camera (Andor, Belfast) and an X-Cite Xylis LED illuminator (Excelitas Technologies, Waltham, MA) fluorescent microscope to visualize vertebrae, and fine forceps, 3 vertebrae were carefully extirpated at 10-15 myomeres posterior to the cloaca. To track the regenerative response, animals were imaged up to 8 weeks after injury using an Axio Zoom.V16 (Zeiss, Jena) equipped with an Orca-Flash4.0 camera (Hamamatsu Photonics, Hamamatsu) and an X-Cite Xylis LED illuminator (Excelitas Technologies, Waltham).

### Atomic force microscopy

Regenerated axolotl vertebrae were surgically removed on pre-chilled plates. Regenerated tissue was flash-frozen in OCT on dry ice. OCT blocks were trimmed and cryo-sectioned at −10°C at a thickness of 20 μm. Sections were collected on electrocharged slides and stored at −80 °C. Atomic force microscopy (AFM) experiments were conducted on a JPK NanoWizard ULTRA SpeedA (JPK-Bruker) AFM system equipped with an AXIO Observer.D1 (Zeiss) inverted optical microscope. MLCT-O10 C (Bruker) tipless cantilevers, of 0.009 N/m measured spring constant via the thermal noise method^59^, were furnished with a borosilicate glass microsphere (SPI Supplies Uniform Microspheres) 15 μm in diameter as previously described^60^.

Immediately after removing a glass slide from the freezer, a 10-30 μl droplet of PBS was placed on the cryosection to prevent it from drying. Subsequently, a fluid cell was mounted and sealed with a two-component silicone (picodent twinsil, picodent, Germany), allowing all AFM micromechanical analysis to be conducted on samples hydrated in 750 μL PBS. Cantilever sensitivity was calibrated via a contact-based method in PBS, as previously described^60,61^. Deflection (volts) vs. z-displacement (μm) curves were recorded on a clean glass next to the sample, over a 400 nm x 400 nm scan area with 32 px x 32 px resolution. This was repeated two to three times until the deviation from the previous sensitivity was less than 1%.

Force curves were collected in both the rim and core (features), visible under brightfield microscopy. On each of the 15 regions of interest (ROIs), 16 force curves were recorded in force volume map mode within a 2 μm x 2 μm scan area at 4 px x 4 px resolution. Force curves were recorded at 1.0 nN applied load (relative setpoint) and with a 2-μm z-length. Measurements were conducted in 6 cryosections at 4wpi without BAPN treatment (2 animals), and 4 cryosections at 4wpi with BAPN treatment (3 animals). Force curves were analyzed with the Hertz model for spherical indenters using a custom Matlab program available on Github (https://github.com/Rufman91/ForceMapAnalysis) as previously described (^60^). A Young’s modulus value was obtained for every force curve. Only indentation moduli values resulting from fits with R^2^ > 0.96 were considered for further analysis, and an average Young’s modulus was calculated for each ROI.

### Buckling instability models

Buckling instabilities with well-defined length scales, and with smooth oscillations generically occur under compressive load due to a compromise between two different elastic materials. For the classical setting of a flat sheet of thickness ℎ and Young modulus *E*_*s*_, under compression and tethered to an infinitely large elastic half-plane of Young modulus *E*_*c*_, simple scaling analysis considering the sheet-bending modulus *E*_*s*_ℎ^3^and the surrounding modulus *E*_*c*_ means that the wavelength of the instability must scale as 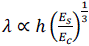, with pre-factors depending on the exact geometry. In the case of a thin shell of thickness ℎ much smaller than its radius of curvature *R*, this limit is expected to hold and predict the buckling instability. More specifically, in our setting, we first only consider peristaltic deformations, i.e. rotationally symmetric, so that the tube displacement can be parametrized by *u*_*r*_(*z*) where r represents the radial direction and z the directional along the tube. We consider a resting state for a tube composed of a stiff shell of thickness ℎ and Young’s modulus *E*_*s*_ as well as a softer core of radius R and Young’s modulus *E*_*c*_. For all materials, we will take the Poisson ratio *ν* to be close to 0.5, i.e. an incompressibility assumption classical for biological materials. Interestingly, neglecting the mechanics of the core reflects a fundamental difference between cylindrical and flat geometries. Computing the elastic energy of the shell from its strain components^62,63^, under the classical Foppl-Von Karman approximation, reveals that whereas compressed flat sheets buckle according to their fundamental mode (i.e. system size), the constraints of cylindrical geometry select a finite length scale for buckling, dependent on the aspect ratio of the shell 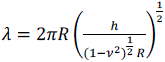. In the regime of a very large radius, we go back to the previous result for flat sheets, dependent on the ratio of stiffness between core and shell 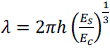 The relevance of each regime depends on the relative mechanics of the shell vs the core, i.e. the dimensionless ratio 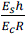. As discussed in the main text, we have computed these numbers via AFM and quantitative geometrical measurements, allowing us to quantitatively assess them. Interestingly, we find an intermediate parameter regime, with a ratio close to 1, as the core is approximately 5 times softer than the shell (*E*_*c*_/*E*_*s*_), but also approximately 4-5 times thicker (*R*/ℎ). We thus compute the wavelength under each regime, at 4wpi, based on 8 independent measurements of the ratio *R*/ℎ, taking in each case the average measured stiffness ratio (*E*_*s*_/*E*_*c*_). Under the geometrical regime, we find *λ* = 505 ± 80 *μm*, while under the differential stiffness regime, we find *λ* = 502 ± 144 *μm*, and thus, interestingly, we are in an intermediate parameter regime where both models make very similar predictions regarding the wavelength. Importantly, reducing the stiffness or thickness of the shell is predicted to decrease the buckling wavelength, again in both regimes of the model, which is in agreement with the results from BAPN treatment. We also remark that in both regimes of the model, the wavelength of patterning can scale with geometric parameters, i.e. as long as *R* and ℎ are larger in larger animals. As such, the wavelength of segmentation can scale with animal size based on purely mechanical considerations. This is very different from predictions of reaction-diffusion models, where additional complex regulatory mechanisms must be invoked to understand how diffusion coefficients or reaction rates change with animal size. Interestingly, we find signatures of this geometrical scaling in the data, as discussed in the main text. In particular, although measuring ℎ systematically across different animal or rod sizes is complicated, as it requires AFM measurements to be precise with respect to the region of higher shell stiffness, we find strong correlation between wavelength *λ* and rod radius *R* in our tail regeneration model, as predicted in the model. In the local injury model, the number of vertebrae *N* should follow from the model *N* = *L*/*λ* where L is the length of the injury. As we predict in our theory that *λ* ∝ 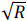, we predict a scaling relationship between *N* and *L*/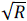, something we observe experimentally, with an increased prefactor in BAPN treatment related to the decrease of *E*_*s*_.

Finally, all models above assume axisymmetric deformations of the rod, i.e. that its overall position remains centered. This also assumes that we can neglect boundary deformations at the ends of the rod, i.e. that they are fixed. These assumptions are valid if the external mechanical constraints acting on the rod are strong. *In vivo*, strong constraints arise from the surrounding mesenchyme. In particular, in the local injury model, the remaining tissues provide strong constraints both at the ends of the regenerating region, and around it. However, in the tail amputation experiments, we hypothesized that external confinement might be much weaker, due to a free boundary condition at the tip of the tail. If we relax the assumption of axisymmetric deformations, and consider a third elastic medium surrounding the rod, of stiffness *E*_*m*_, a second mode of buckling can generically occur, as observed for instance for microtubules under load^62,64^ where the entire rod buckles out of plane. This is exactly what we observe in the 100*μM* BAPN scenario, providing additional confirmation for the hypothesis of pressure-induced buckling of the rod also operating under full tail amputation. In this case, and simplifying the rod material properties to the stiff shell, the wavelength can be written as 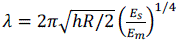. This suggests that for a shell stiffness of around 6kPa, one can explain the global buckling length scale (1-2mm) with realistic stiffnesses of the surrounding mesenchyme of a few hundred Pascals.

### scRNAseq of mature and blastema tissue

Tissue dissociation, and preparation for scRNAseq was performed as previously described^20^. In brief, tail tissue was dissociated in a dissociation solution (0.1 units/μL DNaseI, 35 μg/mL Liberase TM in 0.8X PBS) using fine forceps to continuously tear the tissue apart for approximately 30-40 minutes. Samples were passed over a 70-μm Filcon filter (BD# 340605) and spun down at 2,000 rcf for 3 minutes. Cells were carefully resuspended in 0.8X PBS and flow-sorted with a FACSAria III (BD Biosciences, Franklin Lakes) using an 85-μm nozzle into 200 μL of RLT buffer (Qiagen, RNA mini kit). Single-cell sequencing was done according to manufacturer’s instructions (10x Genomics, Pleasanton, CA), with v3 chemistry used for all experiments except for library 65766, where v2 was used (Extended Data Fig. 8d). We used 5’ 10x sequencing to capture viral-barcode-labeled samples with minor modifications^65^. Ten pmol of a GFP-specific reverse primer and a 10x-specific overhang was spiked into the 5’ 10x droplet capture mix. An aliquot of the cDNA was amplified using a truseq read1 forward primer and a GFP-specific reverse primer with a nextera read2 overhang. This generated amplicons to perform targeted sequencing and was integrated with the data from the 5’ 10x single-cell transcriptome analysis.

Custom 5’ 10x primers used:

5’_10x_GFP_rev_spikein: 5’ AAGCAGTGGTATCAACGCAGAGTACTGGTGCAGATGAACTTCAGGG TC 3’

GFP_Truseq_GFP_fwd: 5’ CTACACGACGCTCTTCCGATCT 3’

GFP_nxt_GFP_rev: 5’

GTCTCGTGGGCTCGGAGATGTGTATAAGAGACAGAACAGCTCCTCGCCCTTG 3’

Viral barcode library production, infection, and clone calling

Viral barcode labeling was performed by modifying existing 3^rd^ generation foamy virus vector systems based on the CellTag principle with modifications^16,18,66^. A 9-nt random barcode sequence and a defined 6-nt tag were cloned shortly behind the TSS of a puc2MD9-SFFV:GFP-WPRE^66^ transfer vector plasmid using Gibson assembly. In brief, SFFV:GFP-WPRE plasmids were amplified in two separate fragments using primers with barcode-containing overhangs, and PCR products were DpnI-treated and purified using para-magnetic beads, and cloned using Gibson assembly. Immediately after assembly, samples were heat-shock-transformed into stellar competent cells (Takara Bio, Kusatsu). SOC outgrowth was performed for 1 hour at 37°C, and a 5-μL aliquot was serially diluted and plated on ampicillin agar plates to estimate cloning efficiencies reaching a minimum complexity of 2*10^6^ colonies. Foamy virus was produced using Lenti-X 293T (Takara Bio, Kusatsu) generated as previously described and frozen at −80 °C for long-term storage^66^. Viral titers were determined by infecting HT-1080 cells with serial dilutions of previously frozen viral preps in multi-well plates. Four days later, plates were imaged using a Cell Discoverer 7 (Zeiss, Jena), and GFP^+^ cells were scored to estimate viral titers. Concentrated virus was injected into either mature or blastema tissue using glass capillary needles and a PV830 picopump (WPI, Sarasota, FL) using standard protocols^66^. We flow-sorted GFP^+^ cells at 21 dpa in two independent experiments and performed scRNAseq using the 5’ 10X platform. Barcoded GFP transcripts were recovered by adapting existing gRNA capture and direct sequencing approaches^65^.

Gibson PCR primers:

CelltagV1_barcode_Fwd: 5’

AGACTGAGTCGCCCGGGTACCGCGGGCCCGNNNNNNNNNACCGGTGGATCCACCGGTCGCCACCATG 3’

CelltagV2_barcode_Fwd: 5’

AGACTGAGTCGCCCGGGTACCGCGGGCCCGNNNNNNNNNCATCACGGATCCACCGGTCGCCACCATG 3’

CelltagV3_barcode_Fwd: 5’

AGACTGAGTCGCCCGGGTACCGCGGGCCCGNNNNNNNNNCGTACAGGATCCACCGGTCGCCACCATG 3’

Celltag_Uni_Rev: 5’ TACCCGGGCGACTCAGTCTGT

AmpR_Fwd: 5’ AATAAACCAGCCAGCCGGAA

AmpR_Rev: 5’ AAGTTGCAGGACCACTTCTG

### Clonal analysis of viral barcode labeling

Non-paraxial mesodermal lineages were excluded from the analysis. Barcodes were extracted and filtered (present in ≥ 2 cells and ≥ 2 UMIs per cell). Clones were called using a Jaccard similarity index of ≥0.45. The axolotl has a measured cell cycle ranging from 53-103 hours^9,67^. Assuming an average cell cycle time of 100 hours, tails infected with V1 4 days prior to amputation would have undergone 5.04 rounds of cell division, tails infected with V2 4 days after amputation would have undergone 4.08 rounds of division, and tails infected V3 7 days after amputation would have undergone 3.36 rounds of cell division. This corresponds to maximum clone sizes of 32.89 (V1), 16.91 (V2), and 10.27 (V3). Reducing the Jaccard similarity index threshold results in unusually large clones being called, which is inconsistent with the known cell-division rate during tail regeneration.

Called clones are displayed in a matrix to visualize clonal diversity (Fig. 3f). Here each row represents a specific cell type, and each column represents a clone. To ensure that clones were not promiscuously called, we computationally combined the two independent replicates, and in this mixed population called clones using a Jaccard similarity index of ≥0.45. We found that 47/48 clones were exclusively made up of cells originating from either replicate 1 or replicate 2, while only a single hybrid clone called was made up of 2 cells from replicate 1 and 1 cell from replicate 2 (Extended Data Fig. 4i). This suggests that with a Jaccard similarity index of ≥0.45 we can reliably call clones in these data-sets and achieve a false discovery rate of 3.6%.

### scRNAseq of embryonic tissue

Axolotl embryos were staged to stage 25-35^68^ and tail buds were dissected in 0.8X PBS under a SZX10 stereo-microscope (Olympus, Tokyo). Individual cells were manually collected using P20 pipettes and prepared for Smart-seq2 using standard protocols in a 96-well format^69^.

### Processing and analysis of scRNAseq data

The quality of the cDNA and resulting sequencing libraries were checked by Bioanalyzer (High Sensitivity DNA Kit, Agilent, 5067-4626). The libraries were sequenced using an Illumina NovaSeq PE150. Sequencing reads were mapped against the axolotl genome^70,71^ using STARsolo (STAR 2.7.5a)^72^, with parameters mimicking Cell Ranger’s (10x Genomics) transcript-counting strategy. Fluorescent marker genes were manually added to the references. Data analysis was performed primarily via Seurat v3.1^73^. Ribosomal protein genes and pseudogenes were excluded from downstream analyses. Generally, cells with a very high or a very? low number of transcripts counts, as well as those with a high mitochondrial transcript proportion, were excluded (see exact information in Table 2). When clusters were identified to be still driven only by low RNA counts in a first screening, cells belonging to these clusters were removed. To enrich connective-tissue cells, some contaminating cell types such as epidermal cells were identified and removed after an initial screening using Seurat. Seurat’s built-in scale function was used to regress out differences in nUMI and percent.mt. Scaling and PCA were always performed on all genes. UMAP was applied to the top principal components (PCs) for visualization. Data integration was performed for the blastema and the uninjured data sets using Seurat’s built-in Harmony^74^ function to reduce the confounding effect of different batches and samples. We performed Harmony using default parameters on the library level, 100 input PCs for blastema integration, and 50 input PCs for uninjured integration. The integration of the uninjured and blastema data sets was performed using Harmony on the library level with slightly adapted parameters (PC=50, theta=2, lambda=9).

### Trajectory analysis

To infer directionality of developmental trajectories during regeneration, we used scVelo^75^. To infer the ratio of spliced and unspliced reads using scVelo, the sequencing data were mapped with STARsolo and its built-in ‘Velocyto’ mode^72^ to obtain spliced and unspliced count matrices, respectively. Afterwards, scVelo was used following the standard procedure (https://scvelo.readthedocs.io/en/stable/VelocityBasics) without dynamical mapping. Fifty input PCs were used for the moments calculation to obtain the arrow velocity graph. Velocity diffusion time was calculated and used as a prior for the identification of links and directionality between the clusters by PAGA.^76^ The diffusion pseudotime presented in Figure 4h was calculated using the standard diffusion map algorithm^77^ in order to set the DMS cells manually to 0 and demarcate them as the starting population.

### Statistical analysis

Statistical analysis was performed using a combination of PRISM (version 9.0.0), and R where appropriate. Student’s t-test and ANOVA were performed using PRISM. MZ structural break test was performed in R.

## References

1. Rodrigo Albors, A., et al. Planar cell polarity-mediated induction of neural stem cell expansion during axolotl spinal cord regeneration. eLife 4, e10230 (2015).

2. Cura Costa, E., Otsuki, L., Rodrigo Albors, A., Tanaka, E. M. & Chara, O. Spatiotemporal control of cell cycle acceleration during axolotl spinal cord regeneration. eLife 10, e55665 (2021).

3. Poss, K. D. & Tanaka, E. M. Hallmarks of regeneration. Cell Stem Cell 31, 1244–1261 (2024).

4. Cooke, J. & Zeeman, E. C. A clock and wavefront model for control of the number of repeated structures during animal morphogenesis. J. Theor. Biol. 58, 455–476 (1976).

5. Baker, R. E., Schnell, S. & Maini, P. K. A clock and wavefront mechanism for somite formation. Developmental Biology 293, 116–126 (2006).

6. Teillet, M. A. & Ledouarin, N. M. Consequences of Neural-Tube and Notochord Excision on the Development of the Peripheral Nervous-System in the Chick-Embryo. Developmental Biology 98, 192–211 (1983).

7. Brand-Saberi, B. & Christ, B. 1 Evolution and Development of Distinct Cell Lineages Derived from Somites. in Current Topics in Developmental Biology vol. 48 1–42 (Elsevier, 1999).

8. Echeverri, K., Clarke, J. D. W. & Tanaka, E. M. In Vivo Imaging Indicates Muscle Fiber Dedìerentiation Is a Major Contributor to the Regenerating Tail Blastema. Developmental Biology 236, 151–164 (2001).

9. Vincent, C. D., Rost, F., Masselink, W., Brusch, L. & Tanaka, E. M. Cellular dynamics underlying regeneration of appropriate segment number during axolotl tail regeneration. BMC Dev Biol 15, 48 (2015).

10. Bussen, M. et al. The T-box transcription factor Tbx18 maintains the separation of anterior and posterior somite compartments. Genes & Development 18, 1209–1221 (2004).

11. Criswell, K. E. & Gillis, J. A. Resegmentation is an ancestral feature of the gnathostome vertebral skeleton. bioRxiv 103, 69–22 (2019).

12. Piekarski, N. & Olsson, L. Resegmentation in the Mexican axolotl, Ambystoma mexicanum. J. Morphol. 275, 141–152 (2014).

13. Krol, A. J. et al. Evolutionary plasticity of segmentation clock networks. Development 138, 2783–2792 (2011).

14. Bessho, Y. et al. Dynamic expression and essential functions of Hes7 in somite segmentation. Genes Dev. 15, 2642–2647 (2001).

15. Fujimuro, T. et al. Hes7 3′UTR is required for somite segmentation function. Scientific Reports 4, 6462 (2014).

16. Biddy, B. A. et al. Single-cell mapping of lineage and identity in direct reprogramming. Nature 564, 219–224 (2018).

17. Kong, W. et al. CellTagging: combinatorial indexing to simultaneously map lineage and identity at single-cell resolution. Nat Protoc 1–25 (2020) doi:10.1038/s41596-019-0247-2.

18. Trobridge, G. D. Foamy virus vectors for gene transfer. Expert Opin Biol Ther 9, 1427–36 (2009).

19. García-García, D. et al. The essential role of connective-tissue cells during axolotl limb regeneration. Preprint at 10.1101/2025.03.30.645595 (2025).

20. Gerber, T. et al. Single-cell analysis uncovers convergence of cell identities during axolotl limb regeneration. Science (New York, N.Y.) 362, eaaq0681–13 (2018).

21. Lin, T.-Y. et al. Fibroblast dedifferentiation as a determinant of successful regeneration. Developmental Cell 56, 1541–1551.e6 (2021).

22. Varner, V. D., Gleghorn, J. P., Miller, E., Radisky, D. C. & Nelson, C. M. Mechanically patterning the embryonic airway epithelium. Proc. Natl. Acad. Sci. U.S.A. 112, 9230–9235 (2015).

23. Gill, H. K., et al. The developmental mechanics of divergent buckling patterns in the chick gut. Proc. Natl. Acad. Sci. U.S.A. 121, e2310992121 (2024).

24. Shyer, A. E. et al. Emergent cellular self-organization and mechanosensation initiate follicle pattern in the avian skin. Science 357, 811–815 (2017).

25. Hannezo, E., Prost, J. & Joanny, J.-F. Instabilities of Monolayered Epithelia: Shape and Structure of Villi and Crypts. Phys. Rev. Lett. 107, 078104 (2011).

26. Hannezo, E., Prost, J. & Joanny, J.-F. Theory of epithelial sheet morphology in three dimensions. Proc. Natl. Acad. Sci. U.S.A. 111, 27–32 (2014).

27. Shyer, A. E. et al. Villification: How the Gut Gets Its Villi. Science 342, 212–218 (2013).

28. Santos-Durán, G. N., Cooper, R. L., Jahanbakhsh, E., Timin, G. & Milinkovitch, M. C. Self-organized patterning of crocodile head scales by compressive folding. Nature 637, 375–383 (2025).

29. Wilmarth, K. R. & Froines, J. R. In vitro and in vivo inhibition of lysyl oxidase byaminopropionitriles. Journal of Toxicology and Environmental Health 37, 411–423 (1992).

30 . Makris, E. A., Responte, D. J., Paschos, N. K., Hu, J. C. & Athanasiou, K. A. Developing functional musculoskeletal tissues through hypoxia and lysyl oxidase-induced collagen cross-linking. Proc. Natl. Acad. Sci. U.S.A. 111, (2014).

31. Timoshenko, S., P. & Gere, J., M. Theory of Elastic Stability. (McGraw-Hill, 1961).

32. Euler, L. Methodus inveniendi lineas curvas maximi minimive proprietate gaudentes sive solutio problematis isoperimetrici latissimo sensu accepti. (1744).

33. Lust, K. et al. Single-cell analyses of axolotl telencephalon organization, neurogenesis, and regeneration. Science 377, eabp9262 (2022).

34. Ma, R. C., Jacobs, C. T., Sharma, P., Kocha, K. M. & Huang, P. Stereotypic generation of axial tenocytes from bipartite sclerotome domains in zebrafish. PLoS Genet 14, e1007775 (2018).

35. Sharma, P., Ruel, T. D., Kocha, K. M., Liao, S. & Huang, P. Single cell dynamics of embryonic muscle progenitor cells in zebrafish. Development dev.178400 (2019) doi:10.1242/dev.178400.

36. Nguyen, P. D. et al. Muscle Stem Cells Undergo Extensive Clonal Drift during Tissue Growth via Meox1-Mediated Induction of G2 Cell-Cycle Arrest. Cell Stem Cell 21, 107–119.e6 (2017).

37. Wopat, S. et al. Spine Patterning Is Guided by Segmentation of the Notochord Sheath. CellReports 22, 2026–2038 (2018).

38. Wopat, S. et al. Notochord segmentation in zebrafish controlled by iterative mechanical signaling. Developmental Cell S1534580724002387 (2024) doi:10.1016/j.devcel.2024.04.013.

39. Peskin, B. et al. Notochordal Signals Establish Phylogenetic Identity of the Teleost Spine. Current biology: CB 30, 2805–2814.e3 (2020).

40. Lleras Forero, L., et al. Segmentation of the zebrafish axial skeleton relies on notochord sheath cells and not on the segmentation clock. eLife 7, 15–4 (2018).

41. Verissimo, K. M. et al. Salamander-like tail regeneration in the West African lungfish. Proceedings of the Royal Society B: Biological Sciences 287, 20192939 (2020).

42. Fröbisch, N. B., Bickelmann, C. & Witzmann, F. Early evolution of limb regeneration in tetrapods: evidence from a 300-million-year-old amphibian. Proc Biol Sci 281, 20141550 (2014).

43. Ellis, M. M. The Gymnotid Eels of Tropical America. (Published by the authority of the Board of Trustees of the Carnegie Institute, Pittsburgh:, 1913). doi:10.5962/bhl.title.43182.

44. Vonk, A. C. et al. Single-cell analysis of lizard blastema fibroblasts reveals phagocyte-dependent activation of Hedgehog-responsive chondrogenesis. Nat Commun 14, 4489 (2023).

45. Fisher, R. E. et al. A Histological Comparison of the Original and Regenerated Tail in the Green Anole, *Anolis carolinensis*. The Anatomical Record 295, 1609–1619 (2012).

46. Khattak, S. et al. Germline transgenic methods for tracking cells and testing gene function during regeneration in the axolotl. Stem Cell Reports 1, 90–103 (2013).

47. Kawaguchi, A. et al. Chromatin states at homeoprotein loci distinguish axolotl limb segments prior to regeneration. *bioRxiv* 2022.11.14.516253 (2022) doi:10.1101/2022.11.14.516253.

48. Aulehla, A. et al. A β-catenin gradient links the clock and wavefront systems in mouse embryo segmentation. Nat Cell Biol 10, 186–193 (2008).

49. Fei, J.-F. et al. Èicient gene knockin in axolotl and its use to test the role of satellite cells in limb regeneration. Proceedings of the National Academy of Sciences of the United States of America 114, 12501–12506 (2017).

50. Fei, J. F. et al. Application and optimization of CRISPR-Cas9-mediated genome engineering in axolotl (Ambystoma mexicanum). Nat Protoc 13, 2908–2943 (2018).

51. Kroll, F. et al. A simple and èective F0 knockout method for rapid screening of behaviour and other complex phenotypes. eLife 10, e59683 (2021).

52. Riquelme-Guzmán, C. & Sandoval-Guzmán, T. Methods for Studying Appendicular Skeletal Biology in Axolotls. in Salamanders (eds Seifert, A. W. & Currie, J. D.) vol. 2562 155–163 (Springer US, New York, NY, 2023).

53. Pende, M. et al. A versatile depigmentation, clearing, and labeling method for exploring nervous system diversity. Sci Adv 6, (2020).

54. Kurth, T. Immunocytochemistry of the amphibian embryo--from overview to ultrastructure. Int J Dev Biol 47, 373–383 (2003).

55. Kurth, T. et al. Electron Microscopy of the Amphibian Model Systems Xenopus laevis and Ambystoma mexicanum. in Methods in Cell Biology vol. 96 395–423 (Elsevier, 2010).

56. Wagner, F., et al. Human Photoreceptor Cell Transplants Integrate into Human Retina Organoids. http://biorxiv.org/lookup/doi/10.1101/2022.08.09.500037 (2022) doi:10.1101/2022.08.09.500037.

57. Choi, H. M. T. et al. Third-generation in situ hybridization chain reaction: multiplexed, quantitative, sensitive, versatile, robust. Development 145, dev165753 (2018).

58. Bensimon-Brito, A. et al. Revisiting in vivo staining with alizarin red S - a valuable approach to analyse zebrafish skeletal mineralization during development and regeneration. BMC Dev Biol 16, 2 (2016).

59. Hutter, J. L. & Bechhoefer, J. Calibration of atomic-force microscope tips. Review of Scientific Instruments 64, 1868–1873 (1993).

60. Kain, L. et al. Calibration of colloidal probes with atomic force microscopy for micromechanical assessment. Journal of the Mechanical Behavior of Biomedical Materials 85, 225–236 (2018).

61. Andriotis, O. G. et al. Nanomechanical assessment of human and murine collagen fibrils via atomic force microscopy cantilever-based nanoindentation. Journal of the Mechanical Behavior of Biomedical Materials 39, 9–26 (2014).

62. Hannezo, E., Prost, J. & Joanny, J.-F. Mechanical Instabilities of Biological Tubes. Phys. Rev. Lett. 109, 018101 (2012).

63. Karam, G. N. & Gibson, L. J. Elastic buckling of cylindrical shells with elastic cores—I. Analysis. International Journal of Solids and Structures 32, 1259–1283 (1995).

64. Brangwynne, C. P. et al. Microtubules can bear enhanced compressive loads in living cells because of lateral reinforcement. The Journal of Cell Biology 173, 733–741 (2006).

65. Replogle, J. M. et al. Combinatorial single-cell CRISPR screens by direct guide RNA capture and targeted sequencing. Nat Biotechnol 38, 954–961 (2020).

66. Khattak, S. et al. Foamy virus for èicient gene transfer in regeneration studies. BMC Dev Biol 13, 9 (2013).

67. Wallace, H. & Maden, M. The cell cycle during amphibian limb regeneration. Journal of Cell Science 20, 539–547 (1976).

68. Schreckenberg, G. M. & Jacobson, A. G. Normal stages of development of the axolotl, Ambystoma mexicanum. Developmental Biology 42, 391–399 (1975).

69. Picelli, S. et al. Full-length RNA-seq from single cells using Smart-seq2. Nat Protoc 9, 171–181 (2014).

70. Nowoshilow, S. et al. The axolotl genome and the evolution of key tissue formation regulators. Nature 1–21 (2018) doi:10.1038/nature25458.

71. Schloissnig, S. et al. The giant axolotl genome uncovers the evolution, scaling, and transcriptional control of complex gene loci. Proceedings of the National Academy of Sciences 118, e2017176118 (2021).

72. Dobin, A. et al. STAR: ultrafast universal RNA-seq aligner. Bioinformatics 29, 15–21 (2013).

73. Butler, A., Hoffman, P., Smibert, P., Papalexi, E. & Satija, R. Integrating single-cell transcriptomic data across different conditions, technologies, and species. Nat Biotechnol 36, 411–420 (2018).

74. Korsunsky, I. et al. Fast, sensitive and accurate integration of single-cell data with Harmony. Nat Methods 16, 1289–1296 (2019).

75. Bergen, V., Lange, M., Peidli, S., Wolf, F. A. & Theis, F. J. Generalizing RNA velocity to transient cell states through dynamical modeling. Nat Biotechnol 38, 1408–1414 (2020).

76. Wolf, F. A. et al. PAGA: graph abstraction reconciles clustering with trajectory inference through a topology preserving map of single cells. Genome Biol 20, 59 (2019).

77. Angerer, P. et al. destiny: dìusion maps for large-scale single-cell data in R. Bioinformatics 32, 1241–1243 (2016).

