## Extended Data Figure 1-9 for "Buckling instability underlies vertebral segmentation during axolotl tail regeneration"

Extended Data Figures

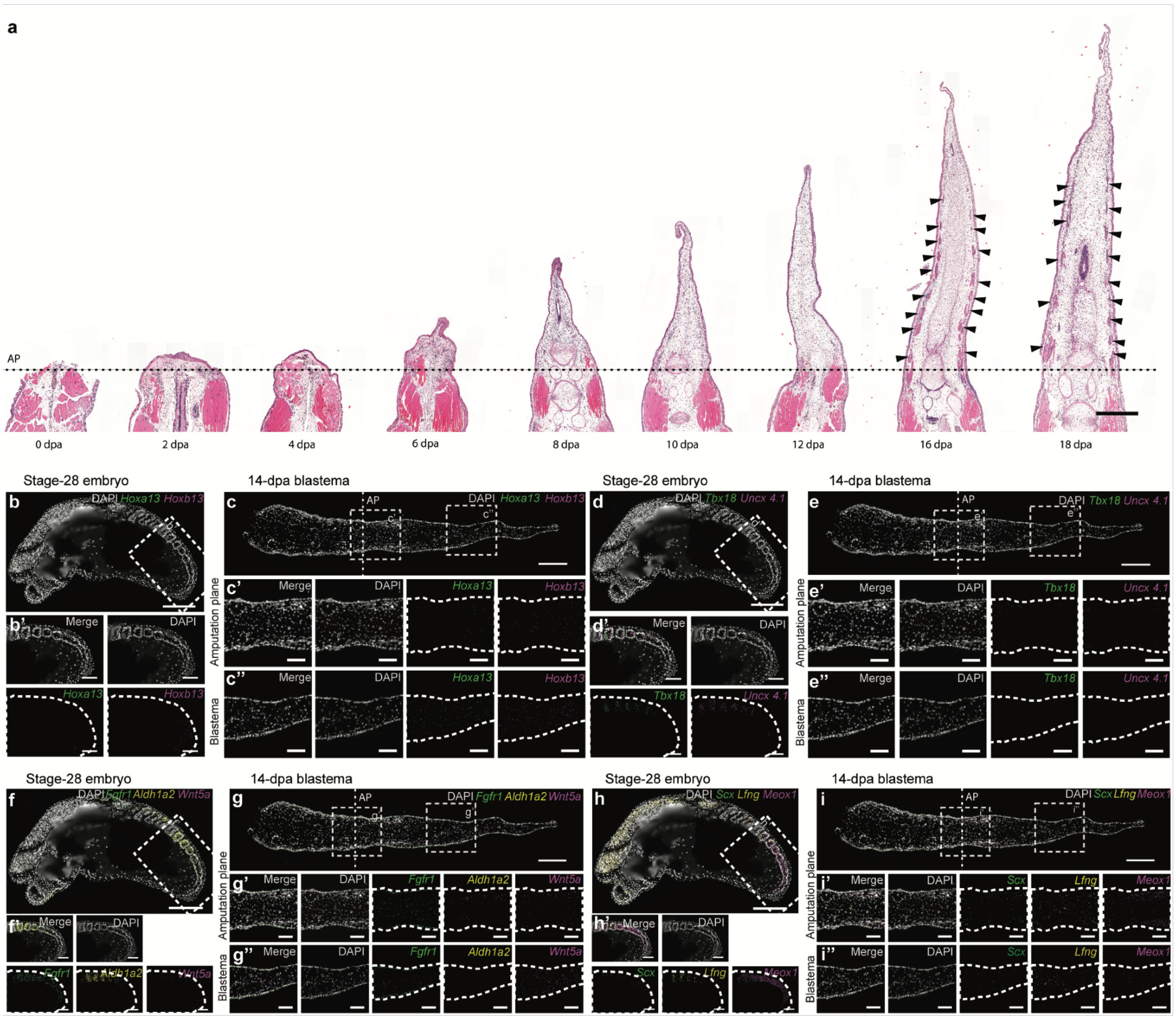


**Extended Data Figure 1: Emergence of segmented myomeres in the absence of hallmarks of somitogenesis during axolotl tail regeneration. a,** Coronal section of H&E-stained tail blastema time series. **b, c,** Representative gene expression patterns for *Hoxa13* and *Hoxb13* in stage-28 embryo (b) and 14-dpa tail blastema (c). **d, e,** Representative gene expression patterns for *Tbx18* and *Uncx4.1* in stage-28 embryo (d) and 14-dpa tail blastema (e). **f, g,** Representative gene expression patterns for *Fgfr1*, *Aldh1a2* and *Wnt5a* in stage-28 embryo (f) and 14-dpa tail blastema (g). **h, i,** Representative gene expression patterns for *Scx, Lfng* and *Meox1* in stage-28 embryo (h) and 14-dpa tail blastema (i). Dashed line indicates amputation plane (AP). Arrow heads indicate myomeres. Scale bars: a-i: 500 µm; b'-h'': 200 µm.

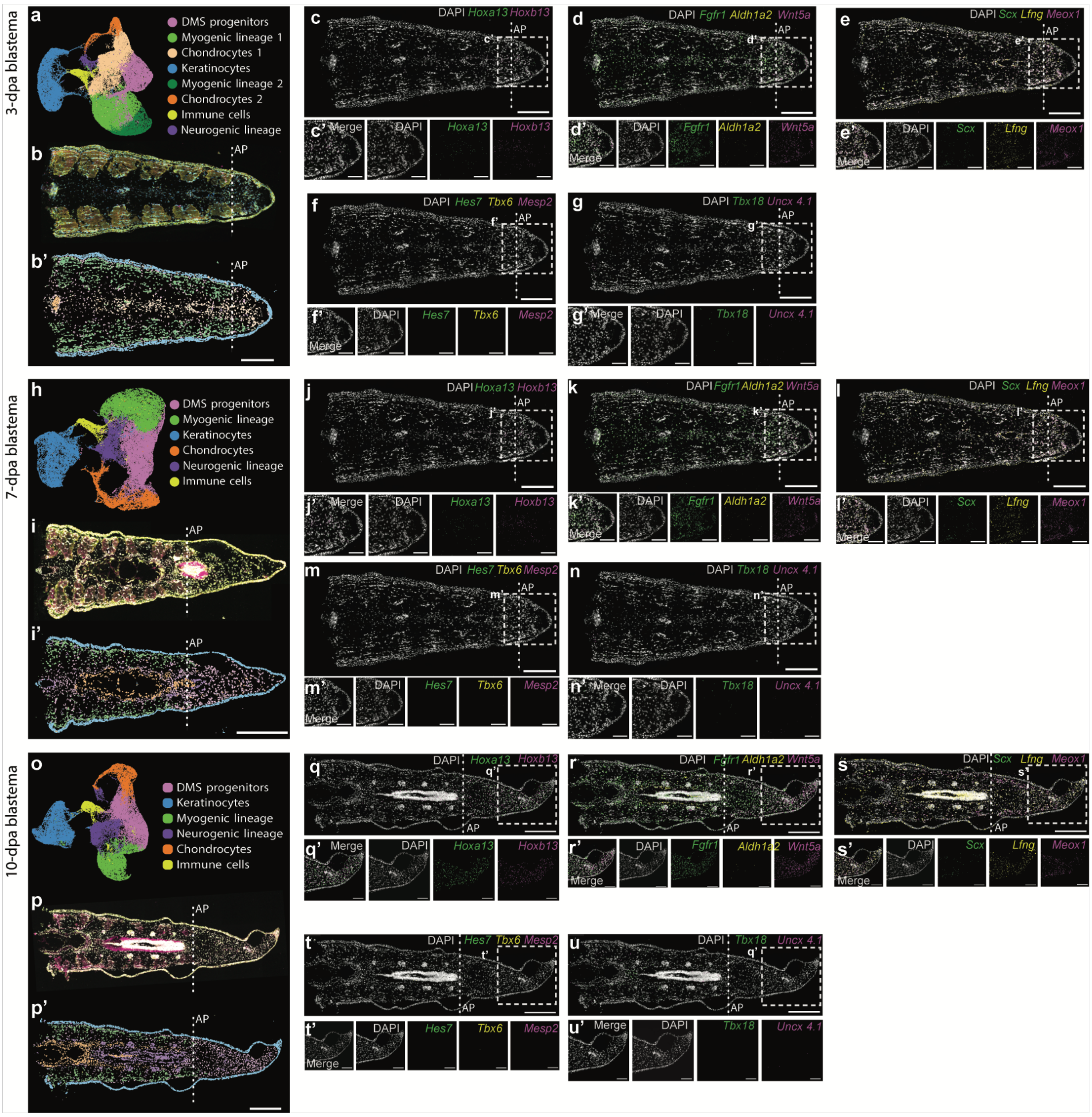


**Extended Data Figure 2: Absence of molecular features associated with somitogenesis in the early tail blastema. a-b', h-i', o-p',** Panels for 3-dpa (a-b'), 7-dpa (h-i'), and 10-dpa (o-p') blastemas, showing UMAP distribution (a,h,o) histology (b,i,p), and spatial distribtion (b',i',p') of identified cell types. **c-f, j-m, q-u,** Xenium expression patterns for selected gene panels, showing the expression patterns for *Hoxa13*, and *Hoxb13* (c,j,q), *Fgfr1, Aldh1a2,* and *Wnt5a* (d,k,r), *Scx, Lfng,* and *Meox1* (e,l,s), *Hes7*, *Tbx6*, and *Mesp2* (f,m,t), *Tbx18,* and *Uncx4.1 (g,n,u)*. Dashed lines indicate amputation plane (AP). Scale bars: b-g, i-m, p-u 500 µm; c'-g',j'-n', q'-u': 200 µm.

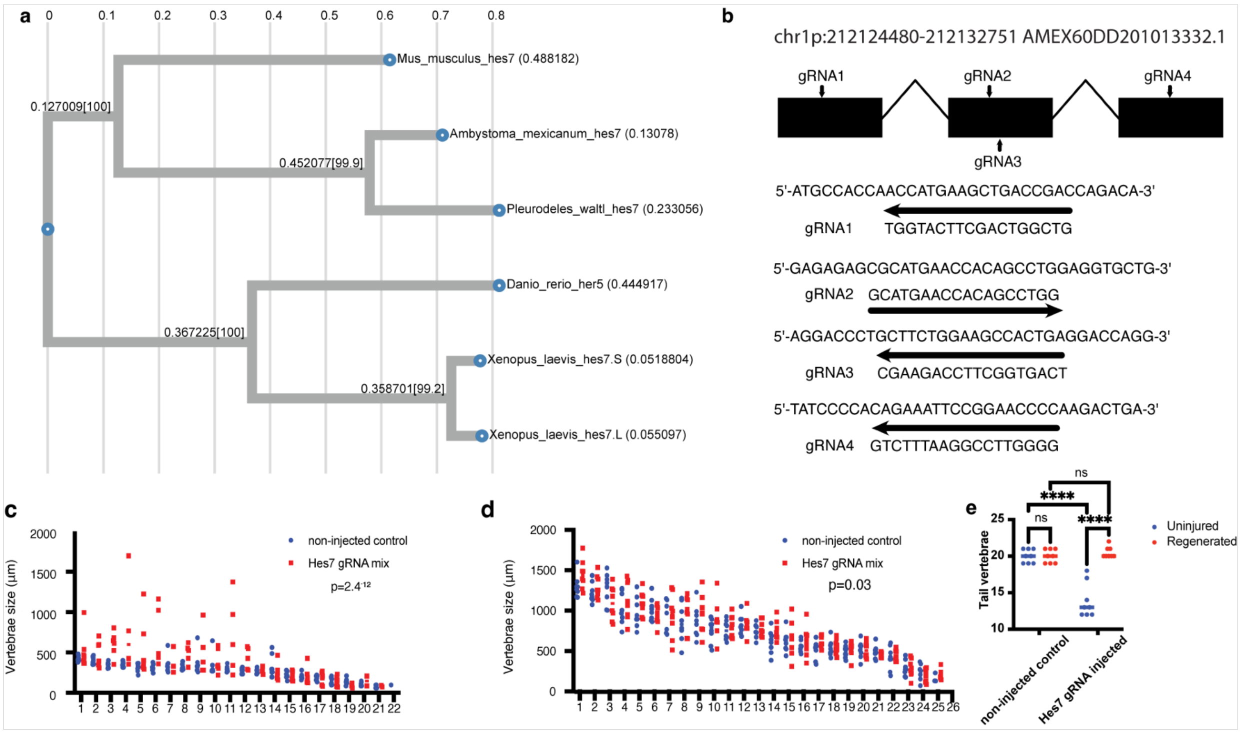


**Extended Data Figure 3: Characterization of *Hes7* crispants. a,** Phylogenetic map showing the relationship of axolotl *Hes7* to several other species including *Mus musculus*, *Pleurodeles waltl*, *Danio rerio*, and *Xenopus laevis*. **b,** Gene model of axolotl *Hes7*, including the locations and exact sequences targeted by gRNA 1-4. **c-d,** Quantification of individual vertebral length from the amputation plane to the tail tip in both uninjured (c) and regenerated (d) animals. **e,** Vertebrae numbers in the non-injected control and *Hes7* gRNA-injected animals in both uninjured and regenerated tails.

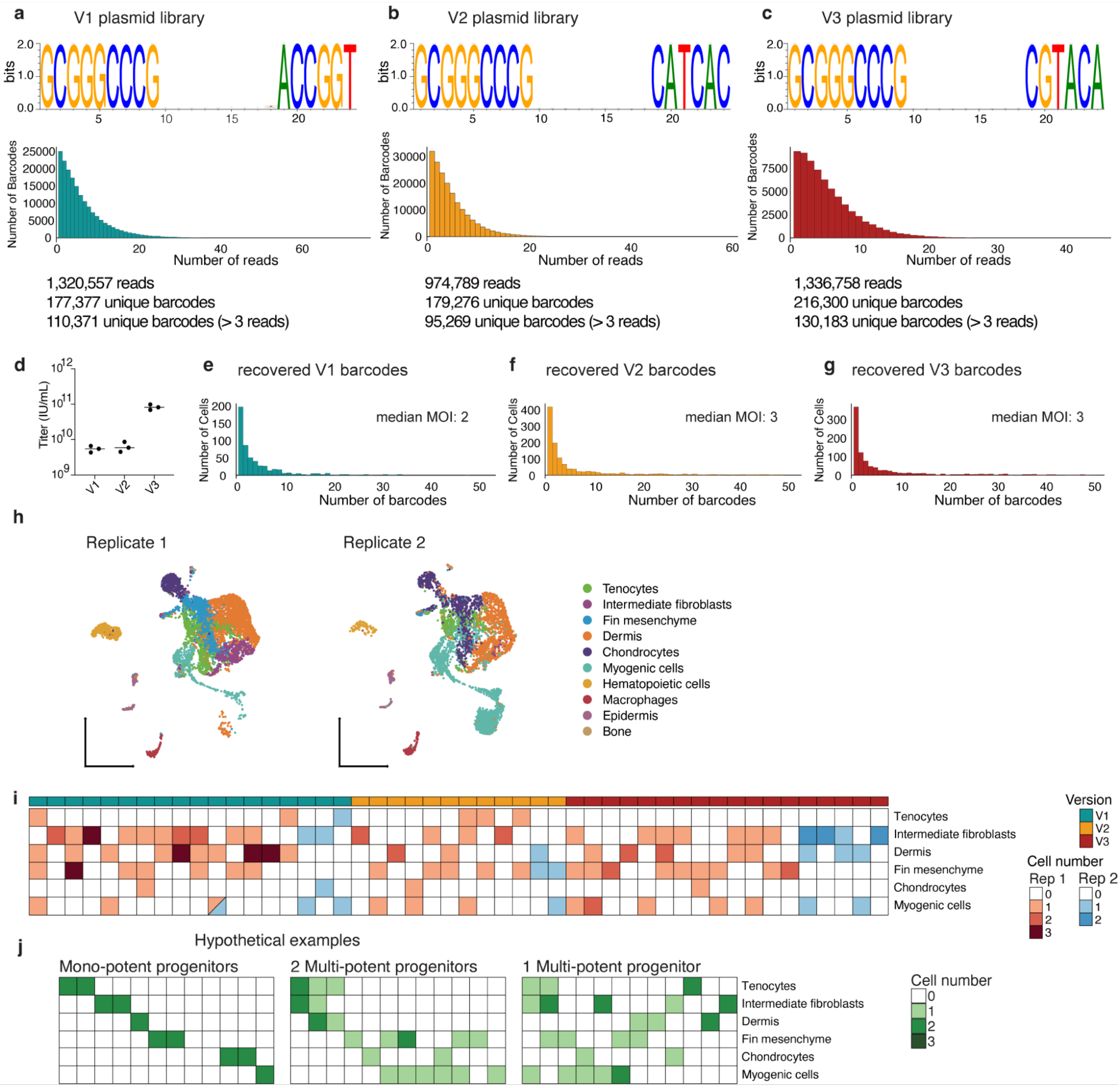


**Extended Data Figure 4: Characterization of foamy virus libraries and barcode-based lineage analysis. a-c,** Sequence logos and barcode representation of V1 (a), V2 (b), and V3 (c) plasmid libraries. **d,** *In vitro* titer assessment of concentrated V1, V2, and V3 viral libraries in HT1080 cells. **e-f,** *In vivo* recovered barcodes per cells from 5’ 10x libraries and experimental MOI assessment of V1 (e), V2 (f), and V3 (g) infected tails. **h,** UMAP representation of the foamy-virus-labeled cells recovered after 5’ 10x sequencing. **i,** Computationally merged matrix-based depiction of identified clones in replicates 1 and 2 with a Jaccard similarity index of ≥0.45 (barcodes present in ≥ 2 cells and ≥ 2 UMIs per cell). Each row represents a cell type.

Each column represents a clone. **j,** Hypothetical examples of lineage-restricted mono-potent clones, two distinct multi-potent progenitors, or a single multi-potent progenitor, in a matrix-based depiction similar to i.

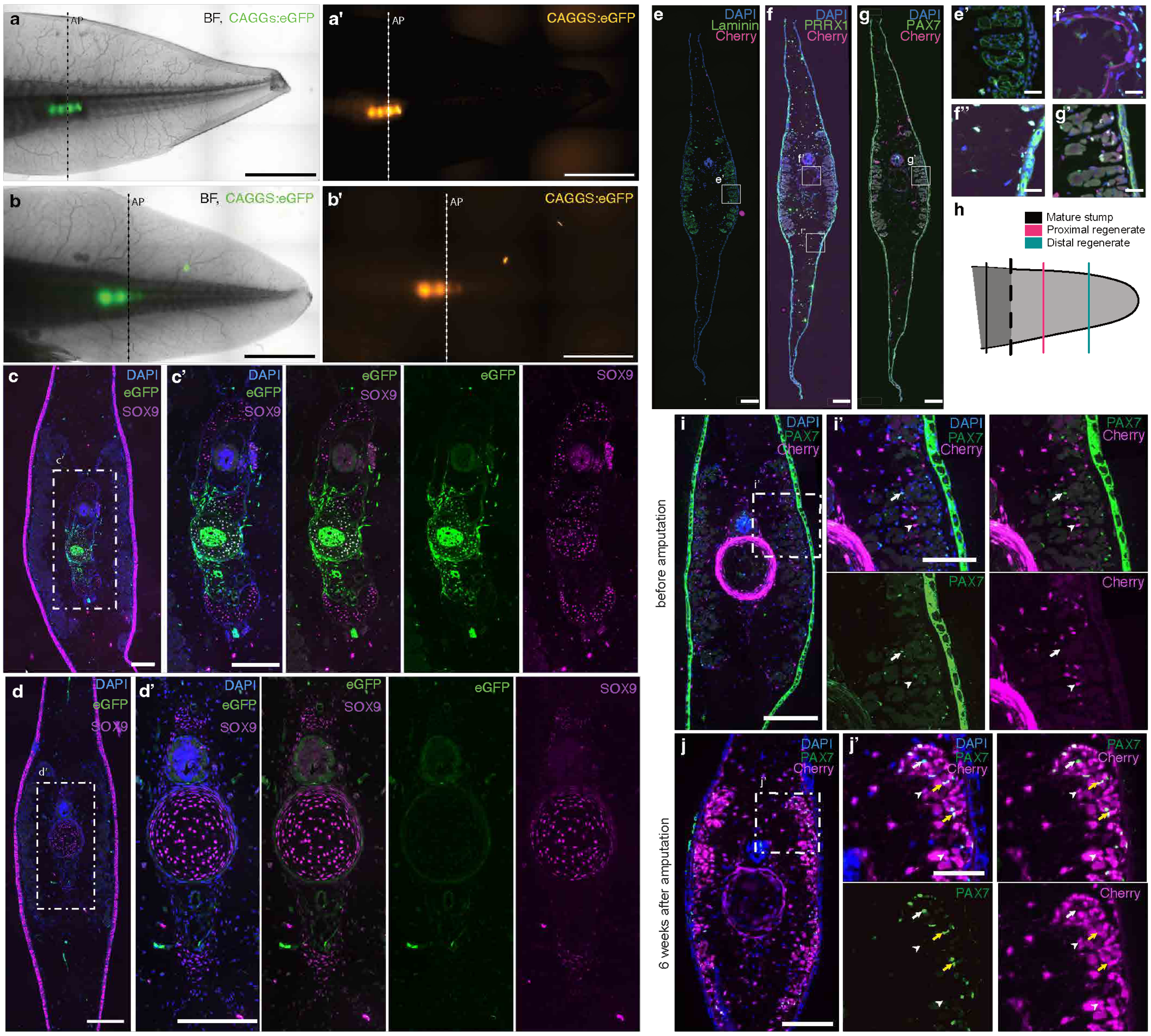


**Extended Data Figure 5: Lineage tracing to identify the cellular contributions to tail regeneration. a-d, Vertebrae grafting and lineage tracing demonstrates a lack of chondrocyte contribution to tail regeneration. a,** *CAGGS:eGFP* vertebrae were grafted into a *d/d* host and amputated after 2 weeks (a' dashed line). **b,** 6-wpa regenerated tail indicating the amputation plane (b' dashed line). **c,d,** IHC staining for GFP (green) and SOX9 (magenta) on a cross section proximal to the amputation plane (c) and on a cross section distal to the amputation plane (d). **c',d'**, enlarged images of bounding box in c, d. **e-h, Sectioning and staining strategy to quantify cellular contribution to tail regeneration. e-g,** Transverse section of the mature tail of the *Col1a2* driver line after tamoxifen conversion, antibody labeled for laminin (e), PRRX1 (f), and PAX7 (g), respectively. **e'-g'**, Zoom images from e-g. **h,**

Schematic diagram of sectioning planes selected for analysis. **i-j, *Col1a2*-labeled progenitors acquire PAX7 identity before differentiating into muscle. i-j,** IHC staining for PAX7 and Cherry on a transverse section of the mature tail of the *Col1a2* driver line after tamoxifen conversion (i) and after regeneration in a 6-wpa tail (j). White arrows indicate PAX7$^+$ Cherry$^-$ cells, white arrowheads indicate PAX7$^-$, Cherry$^+$ cells, yellow arrows indicate PAX7$^+$, Cherry$^+$ cells. Dashed lines indicate amputation plane (AP). Scale bar: a, b: 5 mm. c, d, i, j: 500 µm. e-g: 100 µm. e'-g': 20 µm. i',j': 200 µm.

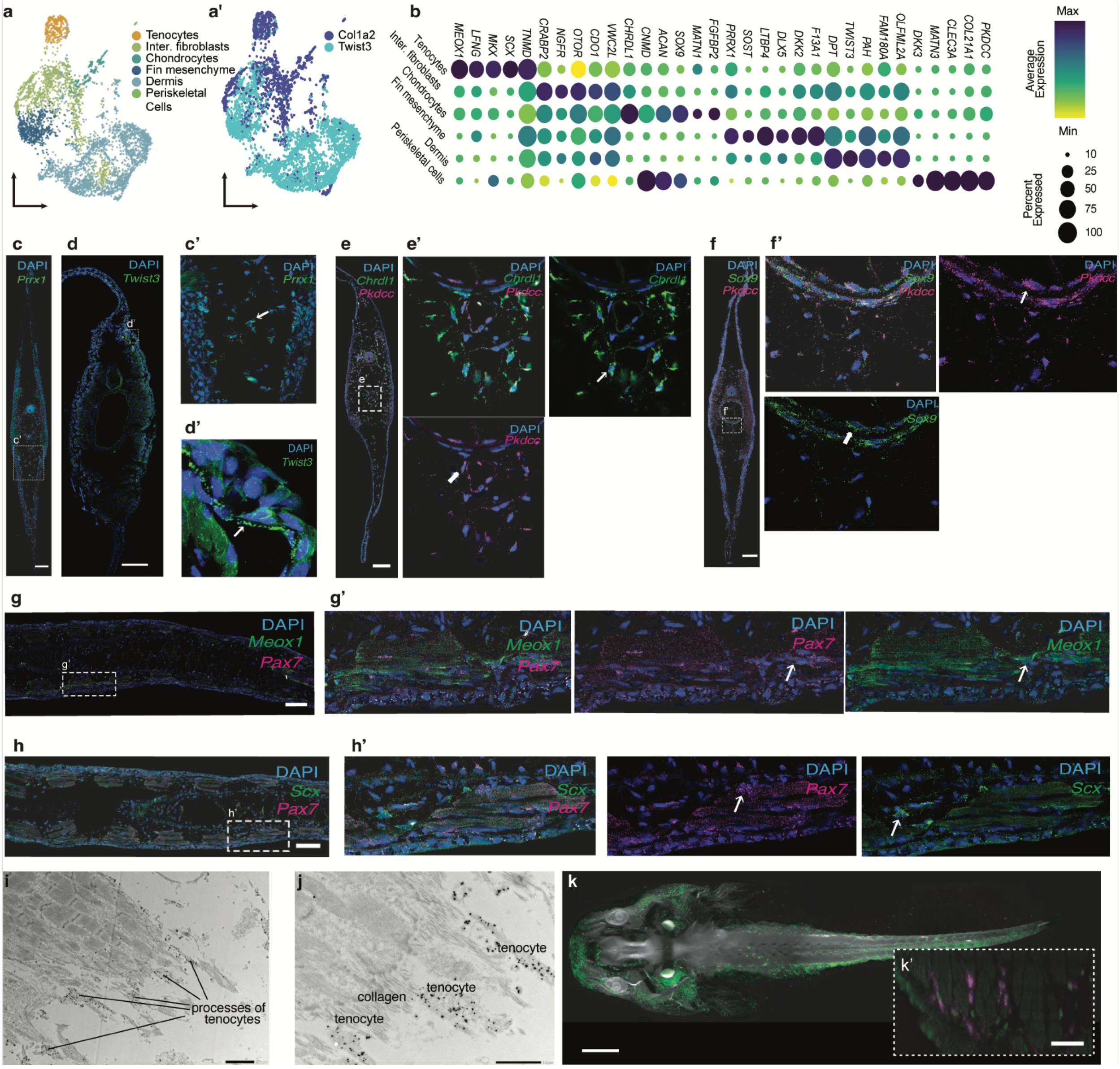


**Extended Data Figure 6: a-b, Characterization of cellular identities in the mature axolotl tail.** UMAP and dotplot representation of different clusters identified in *Col1a2*-, and *Twist3*-converted cells in the mature axolotl tail. **c-f,** HCR-ISH for *Prrx1* (c), *Twist3* (d), *Pkdcc* and *Chrdl1* (e), and *Sox9* and *Pkdcc* (f) in the mature axolotl tail. **g-h,** HCR-ISH for *Pax7* and *Meox1* (g), *Pax7* (g,h), and *Scx* (h) on longitudinal sections of the mature axolotl tail. **i-j,** Immuno-gold anti-RFP labeling on longitudinal sections of the mature *Col1a2*-converted tail. **k,** PCNA labeling on wholemount *Col1a2*-labeled axolotl. **k',** Optical longitudinal section of an axolotl, visualizing both PCNA (green) and Cherry (magenta) labeled cells through *Col1A2* driver. Scale bars: c, h: 200 µm, i: 2 µm, j: 1 µm, k: 2 mm, k': 100 µm

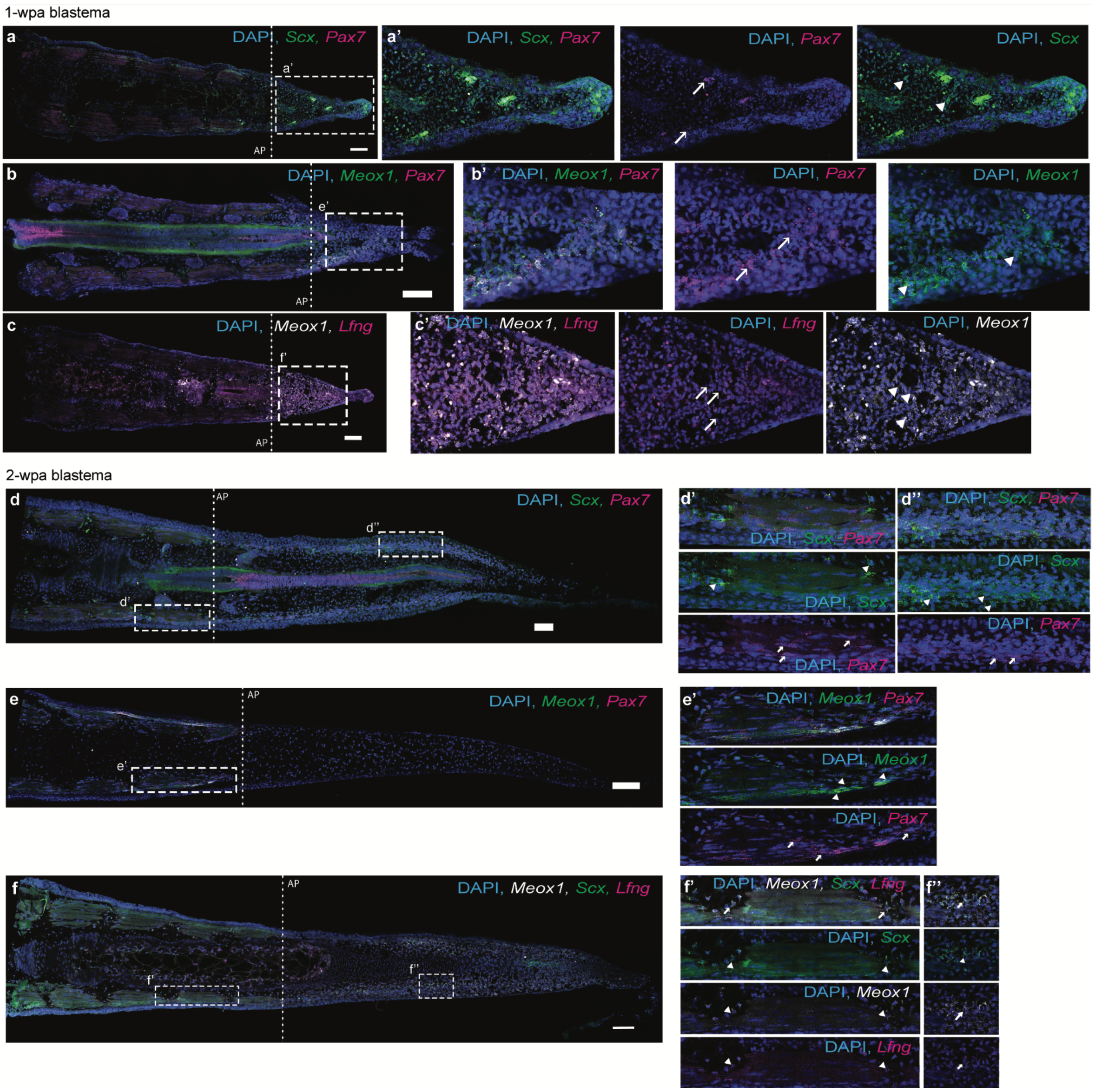


**Extended Data Figure 7: Characterization of cellular identities of the tail blastema**. **a-c,** HCR-ISH on longitudinal sections at 1 wpi for *Pax7* and *Scx* (a), *Pax7* and *Meox1* (b), and *Lfng* and *Meox1* (c), **d-f,** HCR-ISH on longitudinal sections at 2 wpi for *Pax7* and *Scx* (d), *Pax7* and *Meox1* (e), and *Lfng*, *Scx*, and *Meox1* (f). a', d', d'': Arrows indicate *Pax7*, arrowheads indicate *Scx*. b', and e': Arrows indicate *Pax7*, arrowheads indicate *Meox1*. c': Arrowheads indicate *Meox1*, arrows indicate *Lfng*, f', f'': arrows indicate colocalization, arrowheads indicate *Scx, Meox,* and *Lfng* Dashed lines indicate amputation plane (AP). Scale bars: a-f' 200 µm.

a

| Sample library | Timepoint (wpa) | Transgenic line | 10x Chemistry | nUMI maxcutoff | nUMI mincutoff | Percent.mt cutoff |
|---|---|---|---|---|---|---|
| 65766 | 0 | col1 | 3′v2 | 10000 | 5000 | 15 |
| 131286 | 0 | col1 | 3′v3 | 20000 | 1200 | 15 |
| 123453 | 0 | col1 | 3′v3 | 15000 | 500 | 15 |
| 123454 | 0 | twist3 | 3′v3 | 10000 | 1000 | 15 |
| 123456/114342 | 1 | col1 | 3′v3 | 15000 | 3000 | 15 |
| 114343 | 2 | col1 | 3′v3 | 10000 | 1000 | 20 |
| 197397 | 3 | d/d | 5′v2 | 30000 | 5000 | 5 |
| 208683 | 3 | d/d | 5′v2 | 2500 | 4500 | 5 |

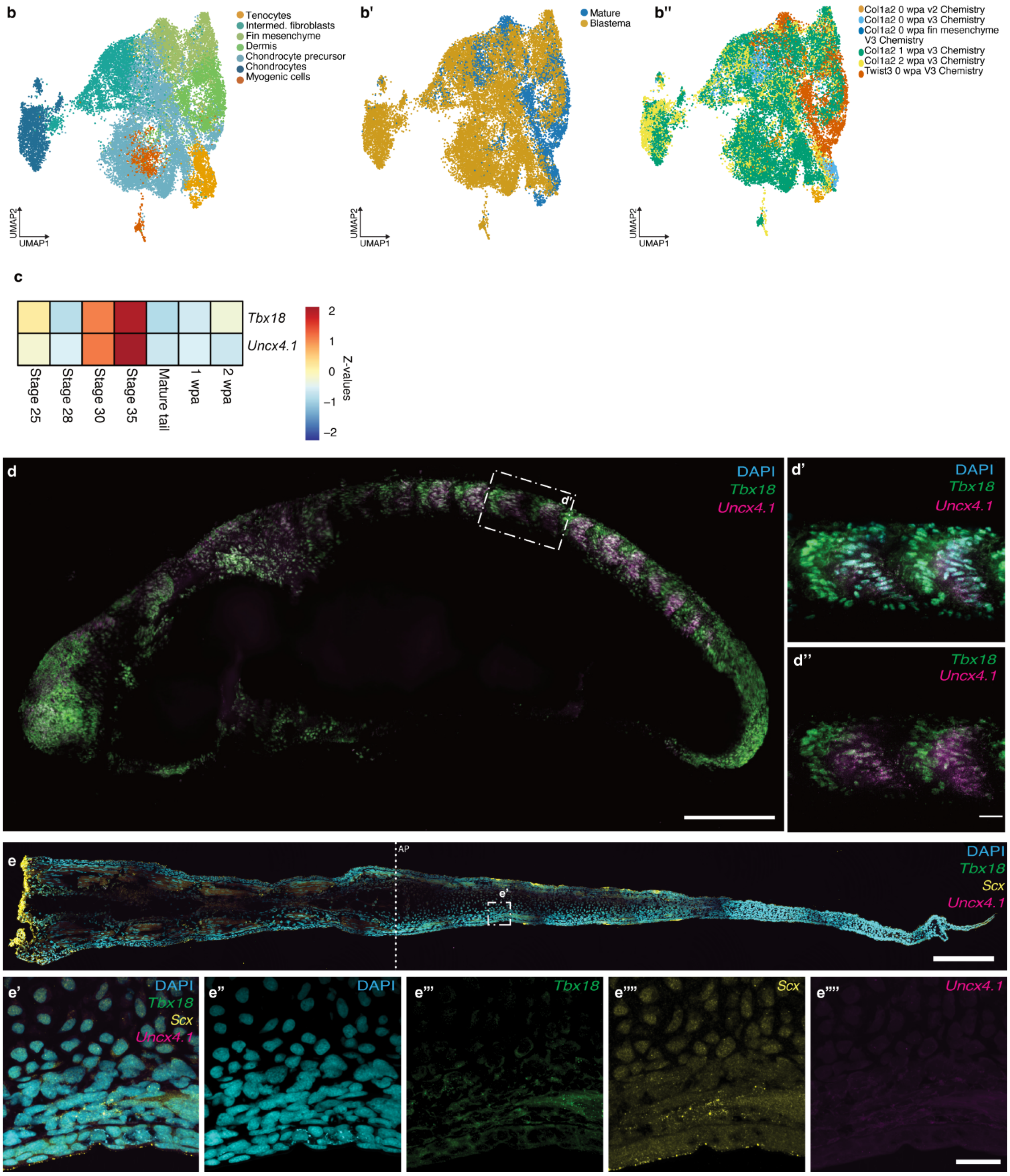


**Extended Data Figure 8. a-b, Integration of mature and blastema single-cell RNAseq datasets. a,** Table showing all single-cell RNA-seq libraries, and corresponding timepoints, transgenic lines and stringency cut-offs for analyses. **b-b”,** UMAP integration of cells from mature and 1- and 2-wpa regenerated tail colored by cell-type identity (b), timepoint (b’), and cell source (b”). **c-e, Lack of resegmentation marker expression during axolotl tail regeneration. c,** Heat-map representation of the pseudo-bulk expression of *Tbx18* and *Uncx4.1* from embryonic and regenerating tail single-cell RNAseq data-sets. **d,** Whole-mount HCR-ISH on a stage-28 axolotl larva for *Tbx18* and *Uncx4.1*, **d’** Bounding box of from d. **d”** Identical to d’ without DAPI. Scale bars: d: 200 µm, d’, d”: 50 µm. **e,** HCR-ISH on a longitudinal section of a 2-wpa tail blastema for *Tbx18, Scx,* and *Uncx4.1*, **e’**, Bounding box from e. **e”**, Bounding box from e, displaying *DAPI.* **e”’**, Bounding box from e, displaying Tbx18. **e””**, Bounding box from e, displaying *Scx*. **e”””**, Bounding box from e, displaying *Uncx4.1*. Dashed lines indicate amputation plane (AP). Scale bars: d: 200 µm, e: 500 µm, d’, d”, e’-e””’: 50 µm.

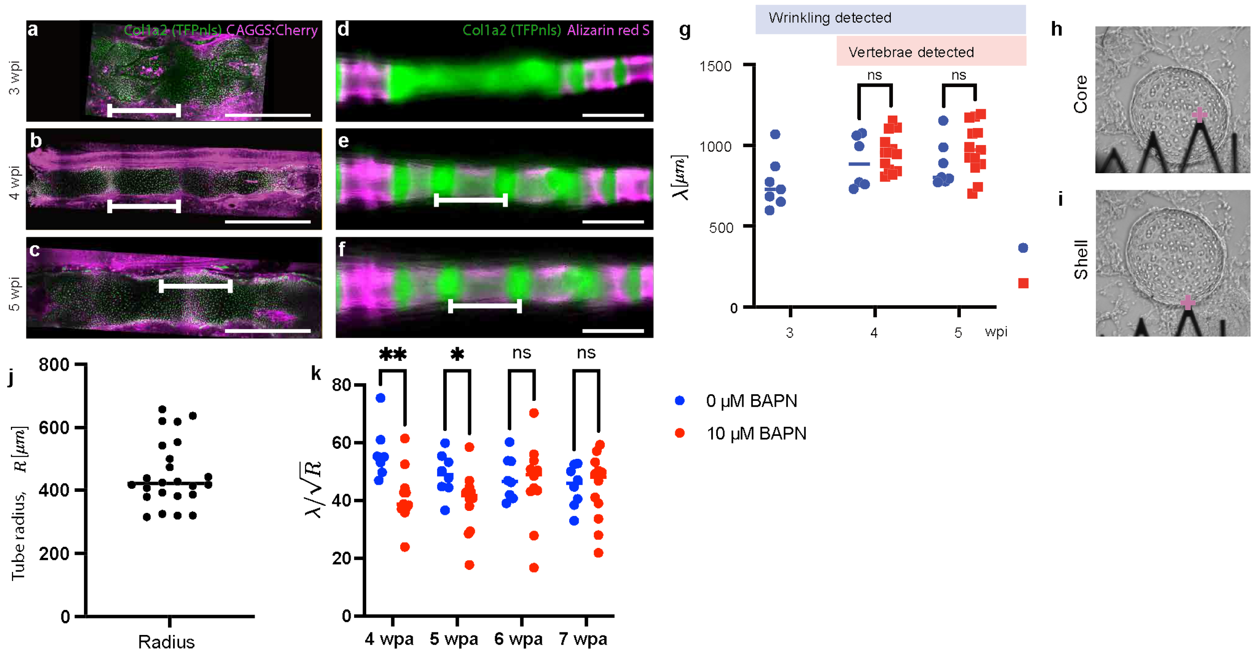


**Extended Data Figure 9: Analysis of the biophysical properties of the regenerating vertebral column. a-f,** Local injury time series of the *Col1a2* driver line crossed to a *CAGGS:Cherry* line (a-c), and the *Col1a2* driver line labeled with Alizarin red S (d-f) from 3-5 wpi. **g,** Quantification of a-f. **h, i,** Brightfield images showing AFM probe location (plus) to record the Young's modulus of the core (h) and shell (i). **j,** Tube-radius quantification based on Fig. 6c. **k,** Quantification of $\lambda/\sqrt{R}$v based on Fig. 6l, m. Scale bar: a-f: 1000 µm.

**Extended Data Table 1:** Gene list and associated information for Xenium spatial transcriptomics.
